# Mesoscale brain dynamics reorganizes and stabilizes during learning

**DOI:** 10.1101/2020.07.08.193334

**Authors:** Yaroslav Sych, Aleksejs Fomins, Leonardo Novelli, Fritjof Helmchen

## Abstract

Adaptive behavior is coordinated by neuronal networks that are distributed across multiple brain regions. How cross-regional interactions reorganize during learning remains elusive. We applied multi-fiber photometry to chronically record simultaneous activity of 12-48 mouse brain regions while mice learned a tactile discrimination task. We found that with learning most regions shifted their peak activity from reward-related action to the reward-predicting stimulus. We corroborated this finding by functional connectivity estimation using transfer entropy, which revealed growth and stabilization of mesoscale networks encompassing basal ganglia, thalamus, cortex, and hippocampus, especially during stimulus presentation. The internal globus pallidus, ventromedial thalamus, and several regions in frontal cortex emerged as hub regions. Our results highlight the cooperative action of distributed brain regions to establish goal-oriented mesoscale network dynamics during learning.

---

Neuronal activity governing behavior is distributed across the brain-wide network of interconnected regions. For example, goal-directed perceptual tasks require computations engaging various regions, including sensory, motor, memory, and reward circuits to generate complex apt behaviors (*1*, *2*). Recent advances in electrophysiological (*3*) and optical methods (*4*) enabled direct measurements of large-scale neuronal activity distributed across multiple regions of the mouse brain in visual (*1*, *5*), tactile (*6*), auditory (*5*, *6*), and olfactory (*7*) discrimination tasks. These studies revealed that activity in many regions represented task-related behavioral variables once expert performance is reached, indicating distributed and parallel sensorimotor processing. Such coordinated network dynamics, appropriate for solving a given task, has to emerge during learning and involves various computational aspects.

Neural ensembles in sensory and higher-order cortical areas learn to identify and discriminate relevant stimulus features in specific contextual settings (*8–11*) and to associate them with particular outcomes such as reward or punishment; thalamus processes and channels parallel streams of sensory (*12*) and motor (*13*, *14*) information to the cortex; and basal ganglia nuclei receive reward prediction error signals (*15*) and transform contextual information into goal-directed behaviors (*16–18*); finally, motor circuits encompassing subcortical and cortical regions are refined to implement the temporally precise execution of apt actions (*19*, *20*). How large-scale brain activity reorganizes during learning, which neural pathways and subnetworks are recruited, and how brain regions dynamically interact to implement the required adaptations remain open questions.

Despite the fundamental importance of learning-related changes in brain activity, it has been difficult to address these questions directly. One experimental opportunity is to apply longitudinal repeated imaging of neuronal activity across the entire training schedule, from naïve to expert (*21*). However, this approach has been applied to either individual (*10*, *22*, *11*) or few regions (*23*) or, in the case of wide-field calcium imaging (*24*), has been limited to the cortical surface. To enable more comprehensive chronic brain activity measurements, we recently introduced high-density multi-fiber photometry (*25*), which can simultaneously target many cortical and subcortical regions of the mouse brain. Here we use this approach to track large-scale brain activity during learning and reveal salient learning-associated changes. Beyond analyzing the changes in individual regions, we apply the framework of functional connectivity (*26*) to study cross-regional interactions among the distributed set of brain regions and to identify salient network properties of emerging functional connectivity.

## Multi-fiber photometry of distributed brain activity during texture discrimination learning

We trained 14 mice in a go/no-go texture discrimination task (*8*). Mice had to lick in response to a ‘go’ texture (rough sandpaper, grit size P100; ‘Hit’ if correct, ‘Miss’ if not licking) and to suppress licking for a ‘no-go’ texture (smooth sandpaper, P1200; ‘correct rejection’, CR, if correct; ‘false alarm’, FA, if licking wrongly; Fig. 1A; Methods). Using an expert criterion of 70% correct performance, mice learned the task within 1-15 days (average 6.2 ± 4.7 sessions; mean ± s.d.; one session per day; one mouse learned the task in the first session; total number of trials ranged from 1999 to 13115). The data were aligned to the first session after reaching expert criterion and divided into a naïve phase (sessions before first expert session) and a subsequent expert phase (Fig. 1B). During task learning, mice established and refined a set of goal-directed actions. Mice developed anticipatory whisking of increased amplitude, likely facilitating early texture discrimination (Fig. 1C; 8.1 ± 2.9° and 15.2 ± 8.7° envelope amplitude in naïve versus expert phase in an early texture presentation window, 3-3.5 s after trial start; p = 0.015, Mann–Whitney U test). Moreover, expert mice reported go-decisions faster than naïve animals (licking onset after texture stop 1.5 ± 0.5 s and 2.1 ± 0.4 s, respectively; p = 0.01, Mann– Whitney U test). In general, behavior became more stereotypical throughout task learning (fig. S1).

**Fig.1.**
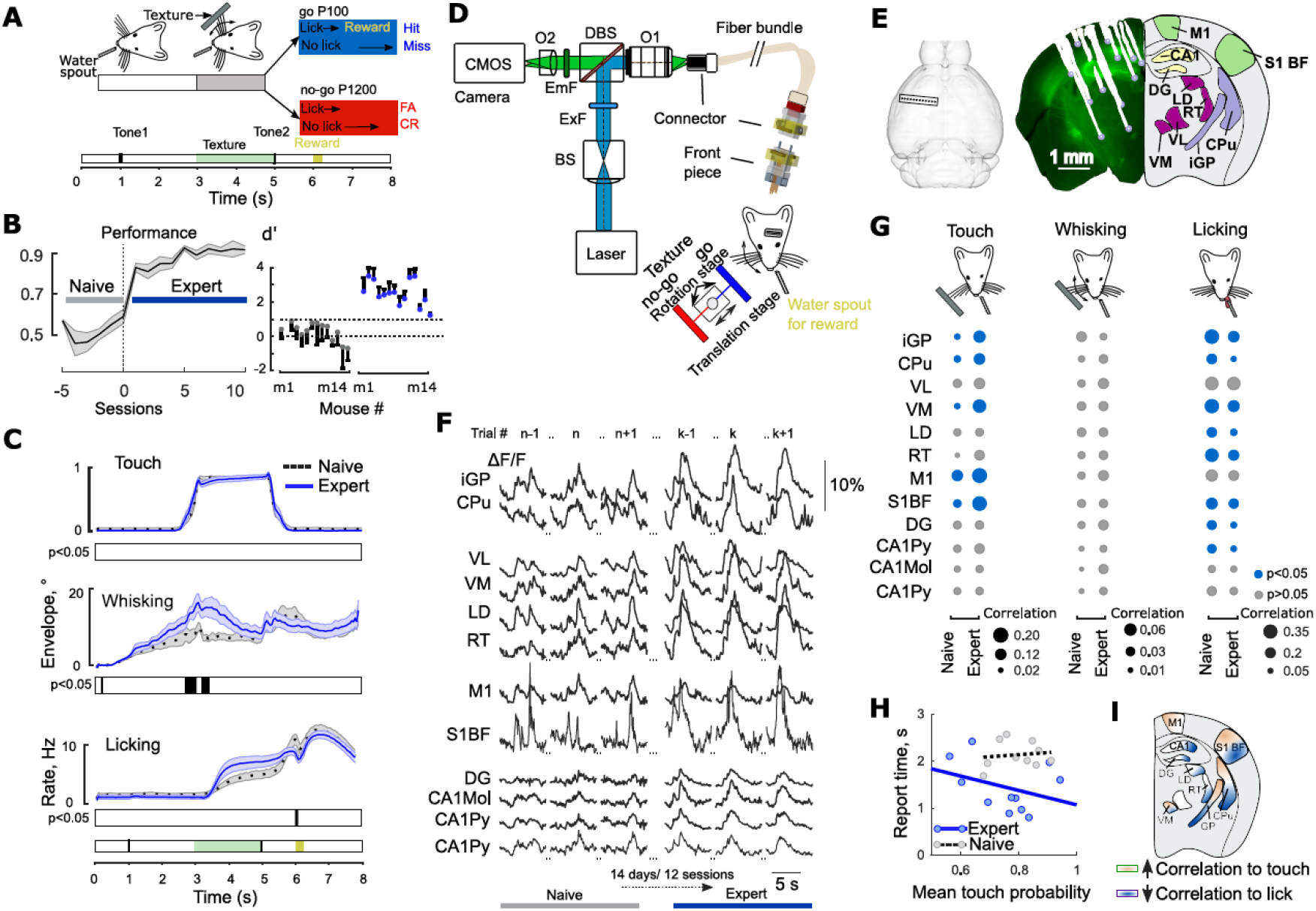
Learning-related changes in behavior and multi-regional brain activity. (**A**) Go/no-go texture discrimination task. (**B**) Left: Aligned task performance averaged across 14 mice (mean ± s.e.m.). Right: d’ for naive and expert phase for each mouse (mean ± s.d.; dashed line, threshold at d’=1.05 [70% correct]). (**C**) Whisker-to-texture touch probability, whisking envelope amplitude and licking rate for naïve (grey, dashed) and expert (blue, solid) mice (n = 14; mean ± s.e.m.; black bars indicate p < 0.01; frame-by-frame Mann–Whitney U test). (**D**) High-density multi-fiber photometry setup. We used 473-nm laser light to excite GCaMP6m and 561-nm light to excite R-CaMP1.07. BS, cylindrical beam shaping lens; O1, objective for coupling the laser beam into the fiber bundle; DBS, dichroic beamsplitter; ExF, EmF, excitation and emission filters; O2, objective forming an image of the fiber array on the CMOS camera sensor. (**E**) Left: Top view of the 12-fiber array position. Right: Reconstructed tracks of the fiber tips (left hemisphere) with corresponding anatomical regions (right; coronal brain section −1.36 mm from bregma). (**F**) Example GCaMP6m ΔF/F traces for the 12 brain regions for naïve and expert phase during Hit trials. (**G**) Pearson correlation coefficients calculated for ΔF/F calcium dynamics and whisker-to-texture touch (for n=11 mice), whisking envelope (for n=11 mice), licking rate for Hit trials (for n=14 mice). Marker size indicates magnitude of correlation coefficient, blue color highlights regions with significant changes from naïve to expert phase (p < 0.05, Mann–Whitney U test). (**H**) Response time (first lick after texture stop) as a function of mean whisker-to-texture touch probability in the 3-3.5 s window (n = 11 mice). Lines are least-squares fit for expert mice (blue solid, R^2^ = 0.12) and naïve mice (grey dashed, R^2^ = 0.015). (**I**) Illustration of brain-wide functional changes for correlations between calcium signals and behavioral variables during learning.

To measure large-scale brain activity during learning, we applied multi-fiber photometry (*25*) (Fig. 1D; 20 Hz frame rate). We chronically implanted multi-fiber arrays targeting 12 regions (n = 10 mice) or 48 regions (n = 4) of the cortico-basal ganglia-thalamic loop and in the hippocampus, combined with viral expression of GCaMP6m or R-CaMP1.07 (Fig. 1E; Methods; for a list of regions and acronyms see Table S1). This approach enables simultaneous recording of bulk calcium signals (expressed as percentage ΔF/F relative to the fluorescence baseline) in all targeted brain regions across many trials from naïve to expert phase (Fig. 1F). For the 12 regions common to all mice, we first analyzed whether calcium signals correlated with behavioral variables and how these correlations might change during learning. Nearly all regional ΔF/F signals showed increased correlation with the texture touch in expert mice, most prominently in cortical S1BF and M1, thalamic VM, and basal ganglia CPu and iGP (Fig. 1G; p ≤ 0.015 for S1BF, M1, VM and CPu; p = 0.03 for iGP; Mann–Whitney U test). Correlation with whisking also tended to increase in most regions, though not reaching significance. Correlation to licking decreased in basal ganglia CPu and iGP, cortical S1BF, thalamic VM, LD, RT, and hippocampal DG, CA1Py regions (p ≤ 0.01 for CPu, LD, DG, CA1Py, p ≤ 0.05 for iGP, VM, RT, S1BF; Mann–Whitney U test). In expert, but not in naïve, mice, higher touch probability also corresponded to earlier licking onsets in Hit trials (Fig. 1H). Overall, the representation of the reward-predicting texture stimulus became stronger during learning, whereas the representation of licking action and reward collection decreased in parallel in several brain regions (Fig. 1I). These results indicate that the mesoscale network spanning cortex, basal ganglia, thalamus and hippocampus underwent a major functional reorganization during learning.

## Widespread shift of activity and discrimination power to the reward-predicting stimulus

We next asked how enhanced behavioral performance is reflected in trial-related activity of different brain regions and their ability to discriminate Hit versus CR trials. In naive mice, there was no significant difference in activity between Hit and CR trials at early trial times—including the initial whisker-texture touch period—but ΔF/F signals diverged later when the animal initiated movements or remained more quiet, refraining from licking (Fig. 2A). The late action-reward related activity was particularly prominent in Hit trials. In experts, the major calcium signal peak in many brain regions had shifted earlier to the texture presentation period (Fig. 2B). In general, two prominent activity peaks were discernible: an early one associated with texture touch and a late one related to reward collection (in addition, several regions showed initial activation upon the auditory trial start cue). To quantify the temporal shift of activity, we calculated the mean ΔF/F signal in an early ‘stimulus window’ (3-3.5 s) and normalized it to the value in a late ‘action-reward window’ (6-6.5 s). The early-to-late ratio significantly increased in 8 out of the 12 regions (Fig. 2C; p < 0.01; Mann–Whitney U test) indicating a widespread shift of neural activity upon learning from the action-reward period to the reward-predicting stimulus (see also Supplementary Notes and fig. S2).

**Fig. 2.**
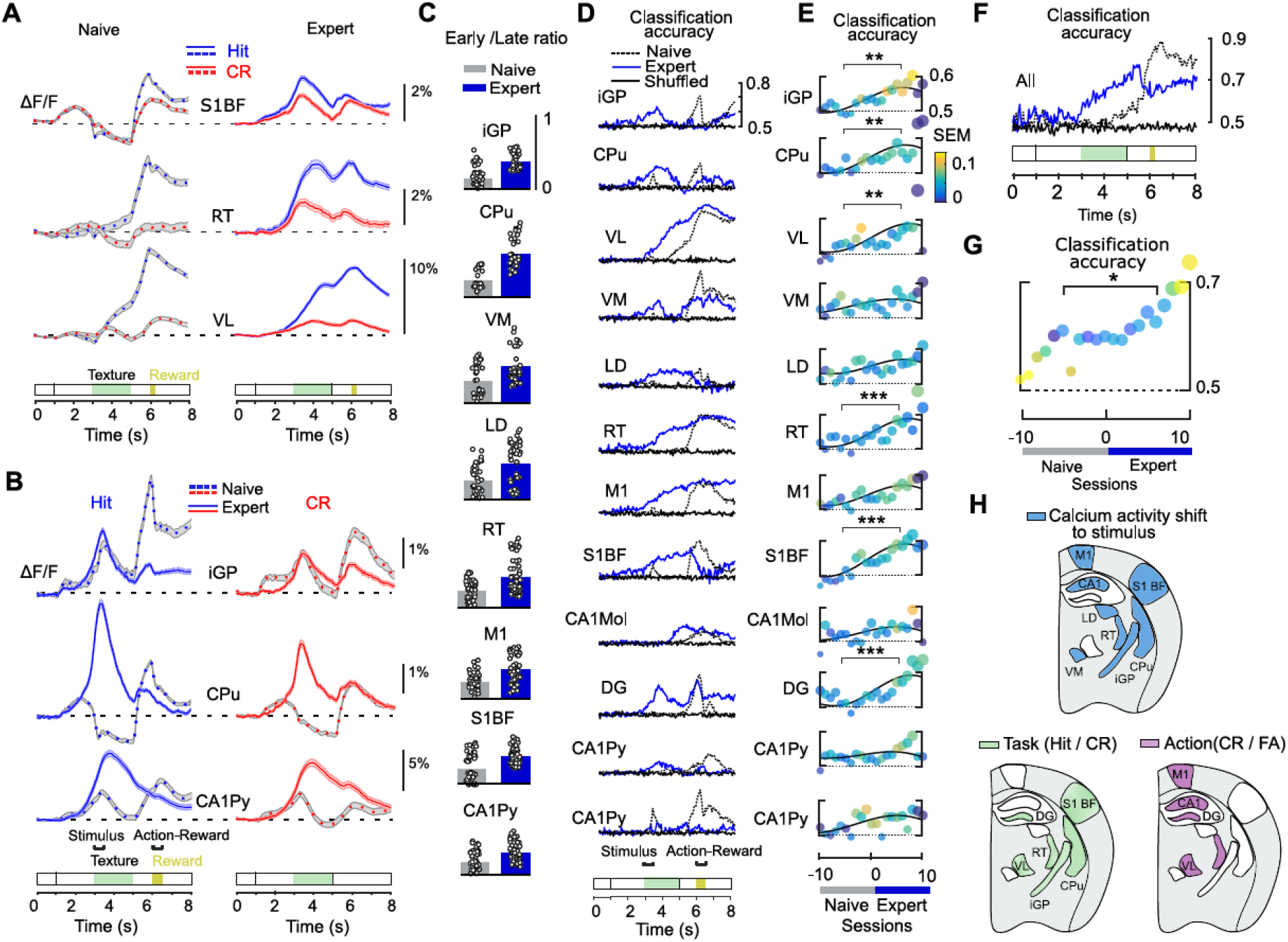
Shift of activity towards reward-predicting stimulus and emerging discrimination power. (**A**) Example of average ΔF/F signals in S1BF, RT and VL from one mouse for Hit trials (blue, n = 324; mean ± s.e.m.) and CR trials (red, n = 326 trials) for naïve (dashed lines) and expert (solid lines) sessions. (**B**) Example of trial-averaged ΔF/F signals in iGP, CPu and CA1Py for naïve and expert sessions. ‘Early’ stimulus window (3 - 3.5s) and ‘Late’ action-reward window (6 - 6.5s) are indicated. (**C**) Early/Late ratio of mean ΔF/F in stimulus and action-reward windows, pooled across naïve and expert sessions (each data point is one session; n = 14 mice). (**D**) Example of the Hit/CR classification accuracy across trial time for naïve (dashed) and expert (blue) trials (same mouse/sessions as in A, B). Classification for shuffled trials labels was at 50% level (solid black lines). (**E**) Hit/CR classification accuracy (n=14 mice, Mann–Whitney U test for naïve vs expert sessions, ±10 sessions around session 0 as illustrated by a grey and a blue bar below. Vertical bars have a scale of 50 to 60% classification accuracy. Color bar shows the s.e.m. across mice. (**F**, **G**) Same as D and E but for the Hit/CR classification accuracy estimated from all brain regions simultaneously. (**H**) Illustration of brain regions shifting peak calcium signals from action-reward to texture presentation period (top, blue) and regions increasing during learning task-related (green) and action-related (purple) classification accuracy.

To assess how well and when regional activities discriminate trial types, we next quantified Hit/CR classification accuracy using linear discriminant analysis (LDA). We first trained the classifier on each trial frame and for each brain region separately (random 80% training data; 20% test data; 20-fold cross-validated). In the naïve phase, classification accuracy reached 60-80% only late in the trial, specifically in the action-reward window (Fig. 2D). In expert mice, classification accuracy increased earlier in essentially all regions and upon texture touch in S1BF, iGP, CPu, and M1. To evaluate stimulus-related classification accuracy across learning while avoiding confounding effects of action-related licking, we trained the LDA classifier on the mean ΔF/F signal during the stimulus window and aligned all sessions to the first expert session. Task-related Hit/CR classification accuracy increased from naive to expert phase in nearly all regions, reaching significance in S1BF, GP, CPu, VL, RT, DG and CA1Mol (Fig. 2E; p = 3e-6, p = 0.0017, p = 0.0021, p = 0.0036, p = 5e-6, p = 6e-6, and p = 0.025, respectively; Mann–Whitney U test; comparing 10 sessions before and after the first expert session). Importantly, improved Hit/CR classification was not solely explained by action-related activity, as indicated by applying the classifier to compare FA/CR (same stimulus) and Hit/FA (same action) trials (Supplementary Notes and fig. S3). Action-related classification accuracy increased during the learning period (defined as ±1 session around session 0) in M1, VL, DG, CA1Py and CA1Mol and gradually decreased in the late expert phase (fig. S3). When we applied LDA across all regions simultaneously, Hit/CR classification reached even higher accuracy (~70%) for late expert sessions (Fig. 2, F and G). Thus, the widespread shift of activity to the stimulus window (Fig. 2H) was paralleled by enhanced task (Hit/CR) classification. Task and action classification improved in distinct but overlapping sets of cortical, basal ganglia, thalamic and hippocampal regions (Fig. 2H). Our observation that Hit/CR classification improved when simultaneously all regions were used for analysis, prompted us to further study learning-related adaptations on the network level by quantifying cross-regional interactions.

## Stimulus-related functional connectivity grows during learning

We examined mesoscale functional connectivity on the two relevant time-scales: throughout learning and within trials. We estimated functional connectivity using transfer entropy (TE) (*27–29*). TE is a directed measure of connectivity, which estimates functional connectivity based on information-theoretic measures of regional activity distributions across trials and time steps. For every pair of regions a functional connection is assigned if a signal of a source region in a past time bin *t*-Δ*t* is predictive of the signal of a target region in the present time bin *t*, but only if the latter cannot be explained by its own past (Fig. 3A; Methods; note that this measure does not distinguish between excitatory and inhibitory action). We used bivariate (considering interactions of region pairs) and multivariate versions of TE (*30*) (the latter finds a connection only if information transfer cannot be explained by any other connection). These approaches provide upper and lower bounds of functional connectivity, respectively (Methods; fig. S4). Using network model with known connectivity, we confirmed low rates of false positives and false negatives (fig. S4 and fig. S5).

**Fig. 3.**
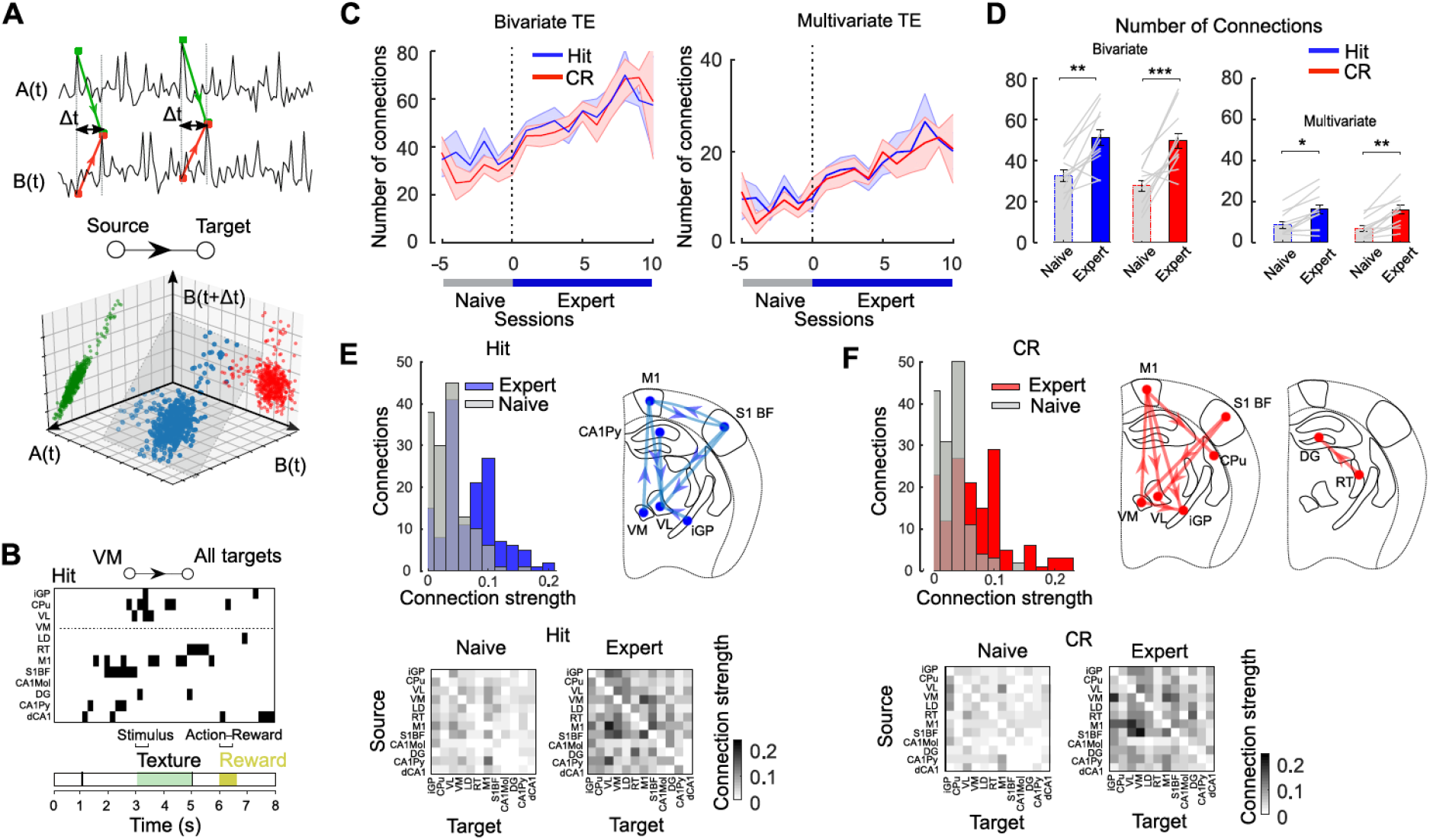
Learning is associated with enhanced network connectivity. (**A**) Illustration example of transfer entropy analysis. Top: The signal B depends on the past dynamics of A. Bottom: a projection of the two time series (blue) onto the subspace A(t)×B(t+Δt) shows a clear correlation (green) whereas projection onto B(t)×B(t+Δt) shows no correlation (red). (**B**) Example of TE estimates across trial time for all connections from VM to other target regions for expert Hit trials (same mouse as Fig. 2, A and B). Black bars indicate frames with significant TE. (**C**) Total number of connections in the stimulus window (3-3.6 s) for Hit (blue) and CR (red) sessions (mean ± s.e.m., n = 13 mice). (**D**) Total number of functional connections for naïve and expert phase, estimated with bivariate (left) and multivariate (right) analysis (mean ± s.e.m., n = 13 mice; gray lines, individual mice; * p < 0.05, ** p < 0.01, *** p < 0.001, Mann–Whitney U test). (**E**) Top: Histogram of connection strength, averaged across mice, during Hit sessions for naïve (gray) and expert (blue) phase. The 8 strongest connections for the expert sessions are shown as a network overlaid with the anatomical position of respective regions. Bottom: matrices of connection strength averaged for naïve and expert phase (averaged over n = 13 mice). (**F**) Same as in E but for CR trials.

We estimated TE in 200-ms time bins to analyze functional connectivity changes across learning in the stimulus window (3.0–3.6 s; 3 time bins; Fig. 3B). Estimates were made separately for each session from all within-session trials. A connection between two regions was labeled as existent if TE was significant in at least one of the stimulus time bins (p < 0.01; compared to shuffled data; Methods). For each session the total number of existing network connections was calculated (the maximum number of possible connections is *k·(k-1)*, i.e. 132 for *k* =12 regions). After aligning sessions to the first expert session and pooling across animals, we found that the total number of stimulus-related connections increases from naïve to expert phase, for both Hit and CR trials and using either bivariate or multivariate analysis (Fig. 3, C and D; in the following we use multivariate analysis as the more conservative approach and show bivariate results in supplementary information). Moreover, we defined ‘connection strength’ as the mean frequency of significant TE bins in a given period (for example, with 3 time bins in the stimulus window, values of 0.0, 0.33, 0.67, and 1.0 are possible for an individual session). The distribution of connection strength (averaged across sessions) shifted towards higher values from naïve to expert phase for both Hit and CR trials (p < 0.001; two-sample Kolmogorov-Smirnov test), with a subset of network connections clearly gaining strengths (Fig. 3, E and F). In expert mice, the 8 strongest connections encompassed cortical, thalamic and hippocampal regions with substantial overlap between Hit and CR trials. We conclude that learning recruits functional connections in the mesoscale brain network in the early stimulus presentation period, consistent with the observed widespread shift of activity towards this trial period.

## Stabilization of functional connectivity during learning

We next asked whether functional connectivity changes occur during specific trial periods and how they compare for different trial types. Expert mice displayed higher functional connectivity primarily during the whisker stimulation period when mice gathered relevant information to discriminate between textures (Fig. 4A). During the later time of the trial, the total number of connections decreased again, converging to naïve level. This observation, together with the finding that the number of connections increases in both Hit and CR expert trials, renders it unlikely that functional connectivity simply relates to animal movements. To assess whether enhanced connectivity may imply that some regions selectively increase their out-going connectivity, we also evaluated changes in the clustering coefficient (*31*, *32*) (Methods) from naïve to expert phase for the stimulus window. The clustering coefficient consistently increased across mice for subcortical regions iGP, VM, and RT in Hit trials, and for CA1Py and RT in CR trials (Fig. 4B; fig. S6 for all regions), indicating that the brain network adapts in a reliable way and stabilizes in an appropriate reorganized state. Consistently, the number of connections shared across consecutive sessions increased from naïve to expert phase (Fig. 4C).

**Fig. 4.**
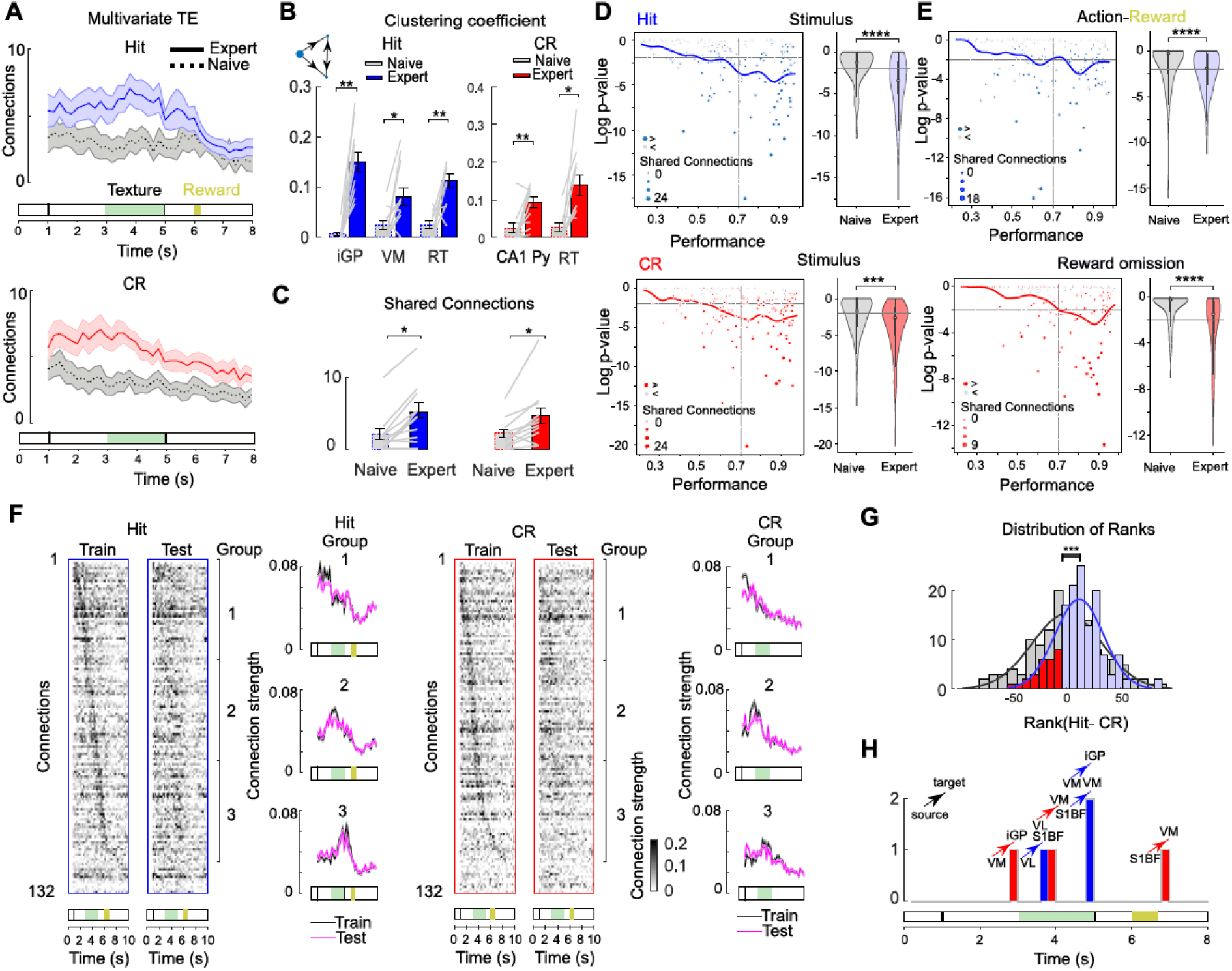
Stabilization of trial-related functional brain network dynamics. (**A**) Total number of connections throughout trial time for Hit (blue) and CR (red) trials (mean ± s.e.m.; n = 13 mice; expert phase: solid lines; naïve phase: dashed line and grey shaded areas). (**B**) Bar plot showing mean ± s.e.m. n=13 mice for the regions which significantly increase their clustering coefficient from naïve to expert performance; Hit and CR expert phase is shown in blue and red respectively; clustering coefficient for the naïve mice is shown in grey. Circles and grey lines show mean values for each mouse. (**C**) Number of connections shared across naïve and expert sessions (mean ± s.e.m. across mice; Mann–Whitney U test). Grey lines indicate the total number of connections for each mouse (n=13 mice with the 12-fiber implants are shown). (**D**) Left: Scatter plot shows the measure of network stabilization (Log p-value) for the observed number of shared connections in the stimulus window versus the task performance. Circles are separate sessions. Solid line is regression to sessions composed of Hit (blue) and CR (red) trials. Dashed horizontal and vertical lines indicate p=0.01 value and 70% performance threshold respectively. Right: Violin plot of Log p-distributions for the naïve and expert phase (Mann–Whitney U test). (**E**) Same as D but for the action-reward period (6 – 6.6 s). (**F**) Strength for all connections across trial time for the expert phase. Data were randomly split into train (n=7 mice) and test set (n=6 mice). Averaged across mice connections for Hit and CR sessions were sorted from early-to late-active according to the peak time after moving average smoothing. Sorting of the train set was also applied to the test dataset. Averaged connection strength for connections 1-40 (top), 41-80 (middle), and 81-120 (bottom) are shown to the right for the train (black) and test (magenta) data. (**G**) Distribution of ranks generated from the difference between Hit and CR connection strength of all connections (positive Hit-dominated ranks, blue; negative CR-dominated ranks, red; null distribution of ranks, grey; Mann–Whitney U test, n = 13 mice). (**H**) Connections which appear more frequently for Hit trials (blue) and CR trials (red) in the expert phase (Wilcoxon signed-rank test p < 0.05; n = 13 mice). * p < 0.05, ** p < 0.01, ***, p < 0.001, **** p < 0.0001.

To further assess network stabilization, we compared session-to-session changes in estimated connectivity with random network dynamics. Specifically, we calculated the likelihood (Log p-value) of obtaining the same number of session-to-session shared connections assuming that the exact connections were randomly permuted for each session (null model). As task performance improved, the likelihood that session-to-session sharing of connections is random markedly decreased in the stimulus window. In the naïve phase, however, network dynamics with random connectivity could explain the underlying number of shared connections (Fig. 4D; pooled across sessions; p < 0.01; Mann– Whitney U test). Session-to-session network dynamics during the action-reward window also showed a stabilization trend (Fig. 4E; p < 0.01; Mann–Whitney U test; bivariate analysis provided consistent results; fig. S7). These results were comparable for Hit and CR trials. We conclude that mesoscale functional connectivity stabilizes throughout learning, especially during the early texture presentation period and with a subset of regions establishing strong network clustering motifs.

We also analyzed trial-related dynamics of connection strength for all possible connections in the 12-region network. For a subset of expert mice (‘train’, n = 7) we sorted the connections according to peak time of their connection strength, revealing a sequential engagement of connections during trial time (Fig. 4F; Methods). To test for consistency across mice, we applied the same sorting to both naïve and expert phase of the remaining ‘test’ subset of mice (n = 6). To visualize the sequential recruitment of connections, we organized connections into three equal groups. The first group (1 - 40) showed high activity early, especially between the auditory cue and texture arrival; second group (41 - 80) displayed highest activity during texture touch, and connections in the third group (81 - 120) were most active during licking and reward time. The group assignment from the train dataset was partially conserved in the test dataset in expert mice (Fig. 4F right, mean ± s.e.m; see fig. S8 for naïve and expert comparison). In experts, approximately one third of connections engaged strongly after the auditory cue and peaked during the texture presentation period, displaying a temporal profile similar to the ΔF/F calcium signals (Fig. 4F; fig. S8 for temporal decomposition). To test for functional connectivity differences between Hit and CR trials, we ranked Hit minus CR strength differences individually for each connection (averaged from 1 – 8 s of trial time) across all mice, summed the ranks over all connections, and compared the resulting test statistic to a null model (Methods). The resulting test statistic was significantly larger than zero, demonstrating that overall connection strength was higher for Hit compared to CR trials (Fig. 4G). To identify differences in the Hit and CR networks, we tested separately for each connection—and in each time frame— whether connection strength differed between Hit and CR trials. Only few connections—all between thalamic regions VM/VL and S1BF or iGP—consistently showed Hit/CR differences across mice at specific trial times (Fig. 4H). We conclude that functional connections are activated in sequential order during trials, with the rough order according to the most salient events preserved across mice. Networks engaged in Hit and CR trials share a substantial amount of connections with few connections showing trial-type specific engagement.

## Internal globus pallidus coordinates functional connectivity in thalamus and cortex

To reveal which regions contribute strongly to functional network formation, we quantified how Hit/CR differences of the clustering coefficient (*31*) emerge during learning. Generally, we found higher clustering coefficients in Hit trials. In naïve mice, Hit/CR differences mainly occurred around the time of reward and they were significant for several regions in the action-reward window (CPu, iGP, VL, S1BF, M1, and CA1Mol; p < 0.05, Mann–Whitney U test; Fig. 5, A and B). In experts, such differences occurred earlier during texture presentation with a significant increase in the stimulus window only for iGP (p = 0.002). The basal ganglia, including its output region iGP, thus may learn to integrate the reward-predicting signal and coordinate appropriate motor commands in the downstream thalamic targets.

**Fig. 5.**
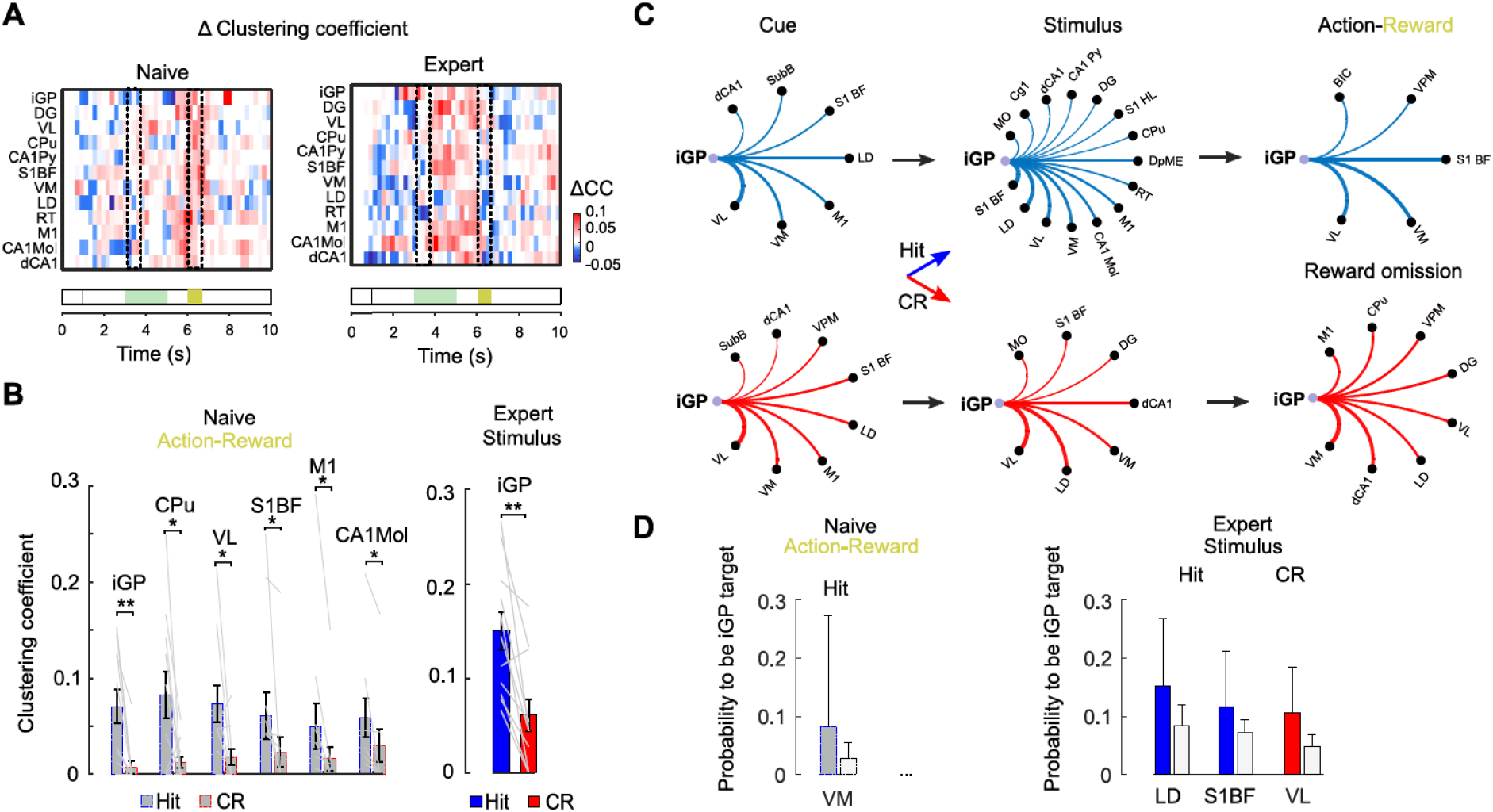
Dynamic changes of regional clustering during learning. (**A**) Hit-CR differences of clustering coefficient (Δ clustering coefficient) for all 12 regions, pooled across mice and plotted versus trial time. Naïve phase, left; expert phase, right. Dashed boxes indicate stimulus and action-reward window. (**B**) Clustering coefficient during Hit and CR trials for the regions showing significant differences in either stimulus or action-reward window (* p<0.05 and ** p<0.01; Mann–Whitney U test; mean ± s.e.m. n = 13 mice; grey lines for individual mice). No significant differences were found in the stimulus window of naïve mice and in the action-reward window for expert mice. (**C**) Regions targeted by iGP during cue (1–1.5 s), stimulus (3–3.5 s) and action-reward (6–6.5 s) windows and in Hit (top row) and CR (bottom row) trials. Line thickness indicates overall number of observed targets pooled across all mice (sorted counter-clockwise from highest to low). (**D**) Probability to be targeted by iGP (mean + 3 s.d., Hit blue and CR red) comparing action-reward period for the naive phase and early stimulus presentation period for expert phase. Grey bars show mean ± 3 s.d. of probability to be targeted by iGP for shuffled connections. Only regions above shuffle mean + 3 s.d. are shown.

In expert mice, we analyzed further which regions contributed to the formation of triangular network motifs as captured by the iGP clustering coefficient. Most frequently, iGP connected to the thalamic regions VL, VM and LD and to cortical S1BF and M1. The connectivity pattern varied throughout trial time, with VL and VM as most prominent targets before stimulation and during the action-reward window whereas S1BF, LD and, VL were targeted during the stimulus window (Fig. 5C). Compared to random networks, S1BF and LD received above chance iGP connections in Hit trials and VL and LD in CR trials (Fig. 5D; in naïve animals only VM and VPM were above-chance in the action-reward window; see fig. S9 for the same analysis for the other brain regions significantly increasing their clustering coefficient during learning, cf. Fig. 4B). These findings highlight the strong impact of iGP on network dynamics and network adaptation during learning (*33*) and especially the pallidal influence on downstream thalamic regions. While the functional connections to thalamus can be explained by the direct anatomical projections (*34*), connections to S1BF and M1 cortical regions require TE transfer via an intermediate region involving at least two synapses.

## Prefrontal regions integrate input from a wide range of brain regions

In 4 mice we implanted 48-fiber arrays (*25*), which allowed us to investigate learning-related network dynamics on an even broader scale, targeting additionally multiple brain regions in frontal and posterior cortex as well as in the amygdala (Fig. 6A; Table S1). Our recordings from this much larger brain network (2256 possible links) substantiated our main findings regarding recruitment and stabilization of mesoscale functional networks during learning. The total number of functional connections increased from naïve to expert phase in the 48-region network, particularly during the texture presentation period (Fig. 6B). Assessment of network stabilization, by comparing session-to-session dynamics with random networks as in Figure 4, again revealed that the likelihood of random session-to-session variability decreased with increasing task performance. This decrease was significant in the stimulus window for both Hit and CR trials when pooled across all sessions and mice (Fig. 6C; p < 0.001, Mann–Whitney U test). We observed no significant gain in stability in the reward-action window (Fig. 6D), possibly due to the higher variability of motor behavior in this trial period. These results highlight the importance of a brain-wide functional network attaining stable and robust dynamics, specifically during the decisive texture touch period when sensory information is pre-processed and parsed to the motor system to generate appropriate motor commands.

**Fig. 6.**
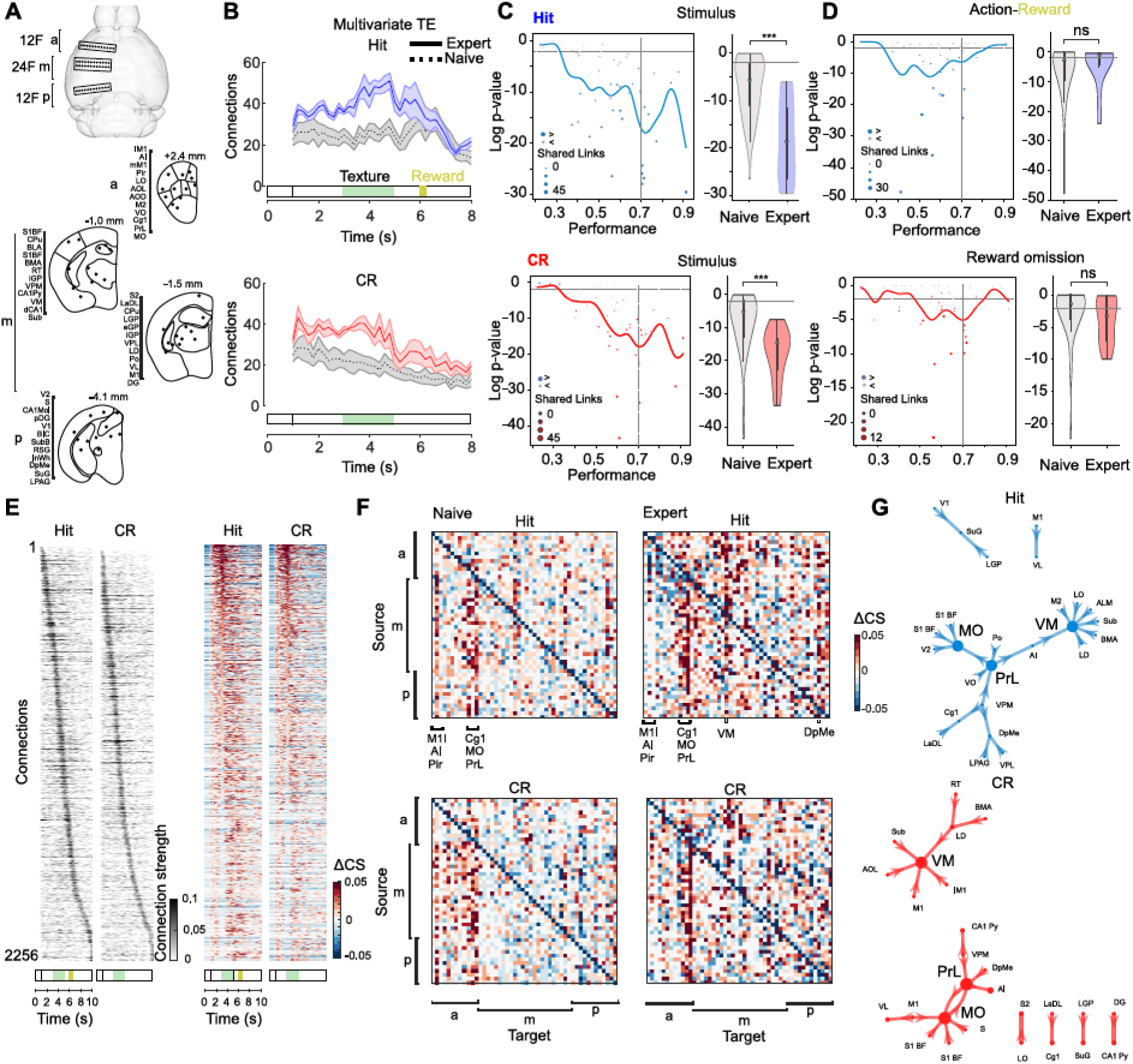
Recruitment of prefrontal cortical circuits during learning. (**A**) Top view of 3 implanted multi-fiber arrays (24-fiber array at 1.4 mm posterior bregma, targeting as a subset the same regions as in 12-fiber experiments; two additional 12-fiber arrays at 2.4 mm anterior and 4.1 mm posterior bregma, respectively). (**B**) Changes in total number of connections throughout trial time (mean ± s.e.m; n = 4 mice for Hit (blue) and CR (red) trials (dashed lines and shaded grey areas for naïve phase). (**C**) Left: Scatter plot shows the measure of network stabilization (Log p-value) in the stimulus window versus the task performance. Circles are separate sessions. Solid lines are regression to sessions composed of Hit (blue) and CR (red) trials. Right: Violin plots of Log p-distributions for the naïve and expert phase (Mann–Whitney U test, ***p<0.001). (**D**) Same as C but for the lick-reward period. (**E**) Mean connection strength across mice (n = 4) plotted during trial time for expert phase for Hit and CR trials. Left: connections were sorted from early-to late-active according to peak time. Right heatmaps: change of connection strength (ΔCS) sorted in descending order according to mean value in the stimulus window. (**F**) Mean connectivity matrices of ΔCS during the stimulus-window. (**G**) Network of regions for the top 1% connection strength for Hit trials (blue) and CR trials (red) in the expert phase.

We further aimed to identify connections with clear changes in connection strength during learning. Connection strength dynamically changed during the trial time (Fig. 6E). As in Fig. 4F connections displayed sequential activation throughout the trial when sorted according to their peak activation times. We also calculated the connection strength change ΔCS in the stimulus window relative to the pre-stimulus period (1 – 1.5 s). About a third of connections increased their strength compared to baseline whereas another third displayed suppression. To identify which connections adapted their strength during the stimulus window, we plotted the mean ΔCS values for the stimulus window as matrices for naïve and expert phase (Fig. 6F). In the naïve phase, regions in frontal cortex (e.g. PrL, Cg1, MO, AI) received strong input from the regions of the basal ganglia-thalamo-cortical network (labeled as m-subsection on Fig. 6A). The subcortical deep mesencephalic nucleus (DpMe) and thalamic VM also received broad input. In the expert phase, Cg1, MO, PrL, DpMe and VM maintained their strong input connectivity acting as integrative hub regions (Fig. 6F and fig. S10), whereas agranular insular cortex (AI) instead received reduced number of inputs. Other regions (e.g. S2) also showed marked changes in their input patterns. These findings demonstrate the global reorganization of the stimulus-associated mesoscale functional connectivity during learning, comprising both strengthening and weakening of specific connections.

To visualize the most salient stimulus-associated connectivity motifs in experts, we plotted the strongest 1% of connections in the stimulus window (Fig. 6G). Prelimbic cortex (PrL), medial orbitofrontal cortex (MO), and thalamic VM appeared as integrating hub regions. Particularly, MO received feed-forward inputs from S1BF during Hit trials and from S1BF and M1 during CR trials, corroborating the idea that signals necessary to update the decision strategy should converge to MO (*35*). We observed strong connections between MO and PrL supporting the hypothesis that these regions interact as decision centers, possibly containing a map of learned stimulus-value associations (*36*). Taken together these results demonstrate that in addition to the basal ganglia-thalamo-cortical loop mice also recruited prefrontal cortical circuits for task learning (*37*).

## DISCUSSION

Multi-fiber photometry provides novel opportunities to study distributed brain activity and its relationship to behavior (*38*, *25*). Here, we demonstrated its chronic application to reveal functional reorganization of brain networks associated with learning. We found that learning-related changes are widely distributed across brain regions. A large fraction of connections in the sub-networks that we sampled with either 12- and 48-fiber arrays were functionally recruited into the task-related mesoscale network. Upon learning, network connectivity increased, starting from the initial cue signaling the texture approach, up to and throughout the decision-relevant texture presentation period. Multiple regions, especially in frontal cortex, increased their activity during the texture approach period, suggesting a role in attentive and anticipatory circuit components. Consistent with previous studies, sensorimotor processing in expert mice then engaged functional connections in cortical (*10*, *11*, *42*), striatal (*43*), thalamic (*44*, *45*) and hippocampal (*37*, *46*, *47*) circuits. A salient learning-related characteristic observed in many regions was the temporal shift of activity and discrimination power from the unexpected reward to the reward-predicting stimulus. This shift is reminiscent to the temporal shift of dopaminergic neuron activity in reinforcement learning studies (*48*, *49*). In experts, touch-related activity may not only reflect refined sensory representations, particularly for the rewarded stimulus, but may also comprise neural signals involved in task-specific movements (*50*, *51*), initiation or suppression of a learned motor program, or prediction of reward-outcome. The temporal shift from reward period to early texture presentation period was also apparent in local network connectivity of iGP, as estimated by the clustering coefficient. We hypothesize that this dynamic shift might have been brought about by the combination of the reward expectation from the dopamine neurons and local striatal circuits integrating this information with the relevant motor commands. In this scenario, iGP acts as a control region coordinating downstream activity in thalamic and cortical circuits. The dense functional connectivity around the basal ganglia also reflects the anatomical organization of basal ganglia-thalamo-cortical loop with its central aim to integrate sensory information and transform it into a goal-directed motor behavior. By studying functional changes on the mesoscale level, our results provide direct evidence of the recurrent interactions across basal ganglia, thalamus, cortical and hippocampal regions.

Several mechanisms could underlie the recruitment of functional connections and their stabilization during learning, including synaptic plasticity (*52*), modulation of inhibition (*53*), cross-regional synchronization (*54*, *55*) that could facilitate recurrent interactions, computations of prediction error from midbrain nuclei (*48*, *49*, *56*, *57*), and common top-down feedback from prefrontal cortical regions (*37*). Resolving the mechanisms underlying the temporal shift of widespread activity from reward to reward-predicting stimulus, will be particularly important, given its fundamental role for credit assignment in reinforcement learning. Chemogenetic or optogenetic manipulations of specific pathways or subsets of pathways could help to verify their causal role in learning-related brain-wide network adaptations. Further, several broad implications can be drawn from the network transformation we observed during learning. For example, increased functional connectivity implies shorter path lengths to connect any pair of regions, facilitating the information flow across the entire network (*26*). Additionally, as networks become more robust, deletion or perturbation of single regions may be compensated by alternative pathways sustaining a large network component (*58*).

In this study we applied two major approaches to analyze learning related changes in the brain activity. First, we focused on stimulus-induced activity and its correlations to behavior in each brain region separately. Second, we applied functional connectivity measures to estimate cross-regional functional interactions, an approach previously mainly used for analysis of fMRI data (*39*, *40*) and more recently also of mesoscale calcium imaging data (*41*). Here, we used MTE as a state-of-the-art model-free information-theoretic metric for functional connectivity. MTE accounts for arbitrary nonlinear relationships between multi-regional signals and excludes spurious functional connections by considering the past of the target as well as the potential influence of other regions. Because it assumes optimal information transfer, MTE only retrieves the most prominent connection if several regions redundantly transfer the same information to a target region. Furthermore, all information-theoretic metrics rely on empirical estimation of entropy, requiring sufficient sample sizes for convergence. Although it is difficult to estimate required sample sizes for real brain networks, repeated optical measurements allow us to collect thousands of trials across behavioral states. Different from fMRI studies, which often assess functional connectivity and changes thereof on the time scale of minutes to hours, the superior temporal resolution of our optical approach enabled direct evaluation of dynamic connectivity across the trial time of the tactile discrimination task. In summary, large-scale calcium recordings across brain-wide regional networks open new avenues to reveal the dynamic changes in functional connectivity emerging during learning. Identification of the salient regional interactions and key features of brain adaptations will help to better understand principles of brain dynamics during learning.

## ACKNOWLEDGEMENTS

We thank Hansjörg Kasper, Martin Wieckhorst and Stefan Giger for technical assistance and Philipp Bethge, Asli Ayaz, Valerio Zerbi, and Denis Burdakov for comments on the manuscript. This work was supported by grants to F.H. from the Swiss National Scien**c**e Foundation (31003A_149858 and 310030B_170269) and the European Research Council (ERC Advanced Grant BRAINCOMPATH, project 670757), by a SystemsX.ch Transfer Project 51TP-0_145729 and through a Roche Joint Collaborative Project.

## AUTHOR CONTRIBUTIONS

Y.S. and F.H. designed the experiments, Y.S. conducted the experiments, Y.S., A.F. and L.N. analyzed data, Y.S. and F.H. wrote the paper.

## DECLARATION OF INTERESTS

The authors declare no competing interests.

## MATERIALS AND METHODS

### Animals and surgical procedures

All animal experiment procedures were carried out according to the guidelines of the Veterinary Office of Switzerland and approved by the Cantonal Veterinary Office in Zurich. A total of 20 male C57BL/6 mice (2-6 months age) were used in this study. GCaMP6m or R-CaMP1.07expression was induced by stereotaxic injections of AAV2.9-hSyn-GCaMP6m and AAV1-EF1a-R-CaMP1.07, respectively. GCaMP6m was used for 12-fiber array implants (n = 7) and R-CaMP1.07 for 12-fiber arrays (n = 3) and 48-fiber arrays (n = 4). Multi-fiber implantation was performed in a separate surgical session, typically 2-5 days after virus injections (*62*). Briefly, we anesthetized mice with 2% isoflurane (mixed in pure oxygen) and maintained body temperature at 37°C with a heating pad. To prevent inflammation and for analgesia, we applied Metacam (s.c., 0.1 μl/g bw). After removal of connective tissue, we polished and dried the skull and applied iBond (Kulzer, Total Etch) to ensure best adhesion of the skull to the connective dental cement. To further stabilize the implant we formed a thin ring of Charisma (Kulzer, A1) on the skull rim. Both iBond and Charisma were light-cured. Small slit-like craniotomies were made to allow for virus injections and fiber-array implantations. First, ~120 nl of virus-containing solution were pressure-injected into all areas of interest at a rate of about 20 nl/min. In order to allow for local diffusion and to minimize potential reflux, the glass injection pipette (10-15 μm diameter) was kept in place for 10 min after each injection. Second, the fiber-array front piece(s) was (were) implanted with the help of a stereotaxic manipulator. The 12-fiber array was implanted at 0.2-mm distance from midline, tilted at an angle of 15 degrees. We oriented the fiber array such that the most lateral fiber efficiently targeted CPu (−1.06 mm from bregma) and the most medial fiber targeted hippocampal areas and posterior motor cortex (CA1, DG, M1; −1.46 mm). Prior to fiber implantation we slightly scratched the dura surface. After, we sealed the craniotomy with vaseline, which melts at body temperature and completely covers the craniotomy. For the 48-fiber experiments, a 24-fiber array was implanted similarly and in an equivalent position as in 12-fiber experiments (−1.46 mm from bregma). Two additional 12-fiber arrays were implanted in frontal and posterior regions (+2.4 mm and −4.1 mm from bregma, respectively). We applied dental cement (Tetric EvoFlow A1) on the skull around the implants followed by UV light curing. A light-weight metal head post was additionally cemented to the skull, allowing for head-fixation during the behavioral experiments.

### Training procedures for the texture discrimination task

Mice recovered for 2 weeks after fiber-array implantation. We habituated mice to head-fixation through a series of short-duration head-fixations in the experimental holder. A day after water scheduling started (for 5 days a week), a mouse was placed in the behavioral setup and first trained to lick following texture presentation. During this shaping period, we presented mostly the ‘go’ texture (rough sandpaper, grit size P100). The auto-reward function automatically opened the water valve at 6 s of trial time conditioning mice to lick on the water spout. The trial structure during the shaping period was the same as for the expert performance, i.e. both auditory cues and the texture were presented. After mice robustly licked, we also presented the ‘no-go’ texture (smooth sandpaper, P1200) and trained mice to discriminate between the two texture types. Mice had to lick for the ‘go’ texture (‘Hit’ if correct, ‘Miss’ if not licking) and to suppress licking for the ‘no-go’ texture (‘correct rejection’, CR, if correct; ‘false alarm’, FA, if licking wrongly). We permitted early licks on go trials whereas licking on FA no-go trials was punished with a prolonged white noise sound. The lick detector was reachable throughout the entire session. Textures were presented pseudo-randomly with no more than three consecutive presentations of the same texture type. In each trial, a first auditory cue (2-kHz tone, two 100-ms pulses at an interval of 50 ms) was given at 1 s after trial start, signaling that the texture was starting to move towards the whiskers on the right side of the snout. The whisker-to-texture touch typically occurred between seconds 2 and 3 of trial time before the texture reached its end position at 3 s. The texture stayed in place for 2 s and was then retracted, also indicated by a second auditory cue (4-kHz tone; four 50-ms pulses at 25-ms interval). A small water reward was delivered in Hit trials (total reward during a full expert session amounted to 1.5 - 3 ml). Mice reported their decision by licking. Typically, mice picked a reward between 5.8 – 6.5 s as can be seen in the increased licking rate starting from 6 s (Fig. 1C).

We trained 14 mice. Mice on average learned the task within 6 days from start of discrimination training (6.2 ± 4.7 sessions; mean ± s.d.; 1 session per day; one mouse learned the task in the first session; total number of trials ranged from 1999 to 13115). We aligned all data to the first session after reaching expert criterion (70% performance) and divided them in a naïve phase (sessions before the first expert session) and an expert phase (Fig. 1B). The average number of trials in both naive and expert phase was around 3500 (3419 ± 2950 and 3556 ± 1638 trials, respectively; mean ± s.d.). However, mice had very different learning rates. One mouse learned the task within the first session, whereas the slowest mouse required 15 sessions.

### Tracking mouse behavior throughout learning

Several behavioral variables were measured throughout task learning. Whisker movement and whisker-to-texture touch times were quantified with markerless pose estimation using DeepLabCut (DLC) (*63*). Videos of the mouse’s snout and whiskers were recorded from the top at 40 frames per second with a camera (A504k; Basler) and 940-nm infra-red LED illumination. We trained DLC to track five body features: (1) the nose tip; (2) the base of the presumed C2 whisker and (3) the whisker shaft position 10 mm away from the base; (4) the bottom corner of the approaching texture; and (5) the intersection of the whisker tip and the texture aiming to quantify the first touch moment. Tracked across frames, the extracted coordinates were used only if DLC confidence was high (p ≤ 0.05). If only one frame had a low confidence the missing value was interpolated as the average of the coordinates from the previous and the following frame. Longer periods of low confidence were assigned NaNs. We calculated the whisker angle *θ* from the dot product *w*_1_ · *w*_2_ = ‖*w*_1_‖‖*w*_2_‖*cos θ* of two Euclidean vectors *w*_1_ (vector nose to whisker-base) and *w*_2_ (vector from whisker-base to the 10-mm shaft position). Next, we transformed the time-dependent whisker angle into whisking amplitude envelope by calculating the maximum minus minimum amplitudes in a 250-ms sliding window. Whisker-to-texture touch was expressed as a binary value for every frame during the trial (0 for p > 0.05, and 1 for p ≤ 0.05). The average of this vector across trials represented the touch probability. The touch probability increased as the texture came into reach of the whiskers and was close to 1 when the texture was standing next to the whiskers. Licking was quantified as event rate from the piezoelectric lick sensor. All behavioral variables were resampled to 20 Hz, the rate of photometry recordings. We calculated the Pearson correlation coefficient between calcium signals and behavioral variables as the covariance of two inputs normalized by the product of their standard deviations. Only correlation coefficients with p ≤ 0.05 were used in further comparison for naïve and expert phase (correlation coefficients with p > 0.05 were set to NaN).

### Multi-fiber photometry setup

We used an Omicron LuxX 473-nm laser for excitation of GCaMP6m and a Coherent OBIS LS 561-nm laser to excite R-CaMP1.07. To achieve stable CW operation, lasers were run at 80% of maximal output power. A variable neutral density filter (NDC-25C-4M; Thorlabs) was used to reduce fluorescence excitation power to ~1.3 mW/mm^2^ per fiber channel at the fiber tip. One or two cylindrical lenses shaped the excitation beam in an appropriate illumination pattern at the object plane of the objective. First, we expanded and collimated the circular beam with an achromatic Galilean beam expander (GBE05-A; Thorlabs). Second, to create a line illumination pattern matching the 12-fiber array we used a 75-mm focal length cylindrical lens (LJ1703RM-A; Thorlabs) placed at ~145 mm distance from the objective (TL4x-SAP; Thorlabs). For the 48-fiber array a rectangular illumination pattern was required. For this purpose, we added a cylindrical lens with 150-mm focal length (LJ1629RM-A; Thorlabs) oriented 90° with respect to the 75-mm focal length cylindrical lens. A dichroic beamsplitter (F58-486 dual line, AHF) coupled the excitation light for both wavelengths (473 nm or 561 nm) into the objective (TL4x-SAP, Thorlabs) and transmitted the fluorescence in the emission spectral windows for GCaMP6m and R-CaMP1.07. To separate fluorescence signals from residuals of the excitation light and to minimize auto-fluorescence generated in a broader spectral range we used emission filters (525/50 nm, F37-516, AHF, for GCaMP6m; and 605/70 nm, F47-605, AHF, for R-CaMP1.07) and a triple-line notch filter at 425, 473 and 561 nm (ZET405/473/561 for Omicron LuxX 473nm and Coherent OBIS LS 561nm lasers, or, alternatively, for a combination of ZET405/488 ZET488/561 Chroma Technology Corp. for Coherent OBIS LX 488nm and Coherent OBIS LS 561nm lasers). To image the end face of the fiber array onto the camera we used a 200-mm focal tube lens (Proximity Series InfiniTube) with internal focusing. The image was created at the back focal plane of the tube lens on the CMOS sensor (ORCA Flash4.0, Hamamatsu camera).

Calcium signals were expressed as percentage ΔF/F relative to the fluorescence baseline (average within the 0–1 s period before the first auditory cue).

### Post-hoc immunohistochemistry

We followed protocols for brain slice preparation as previously described (*25*). Mice were anaesthetized (100 mg /kg bw ketamine and 20 mg /kg bw xylazine) and perfused transcardially with 4% paraformaldehyde in phosphate buffer, pH 7.4. After perfusion, tissue was removed from the skull and the head including the multi-fiber implant was additionally fixated in 4% paraformaldehyde for one week. Then, the ventral (bottom) side of the skull bone was removed and the brain was carefully extruded. Coronal sections (75 – 100 μm thickness) were cut with a vibratome (VT100, Leica). Stained sections were mounted onto glass slides and confocal images were acquired with an Olympus FV1000. Sequential imaging of coronal slices allowed us to track fiber shafts and to verify the 3D positions of the fiber tips. The final assignment of each fiber channel was completed after the fiber tip positions were tracked on the histological slices and aligned to the Mouse Brain atlas.

### Functional connectivity estimated by transfer entropy

In order to quantify compute functional connectivity (the amount of cross-regional interaction) we used transfer entropy (TE) as a directed information-theoretic metric for estimating the relationship between the past activity state of a source region and the present activity state of a target region (*27*, *29*, *30*). In its basic form, the Bivariate TE (BTE) between variables *X*_*t*_ and *Y*_*t*_ is defined as the Mutual Information (MI) between the current value of *Y*_*t*_, and the past value of *X*_*t*−1_, conditioned on the immediate past values of *Y*_*t*−1_.

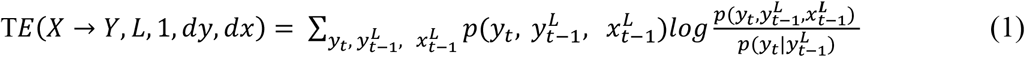

where 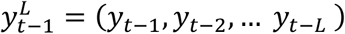 and 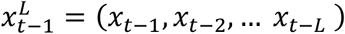. Given the source and target time windows *L*, TE considers the multiple past time-delayed values of the source and target (from *t* − 1 to *t* − *L*) on the TE. It proceeds to select the source time delay value that maximizes the TE. We will refer to this value as the time lag. It approximates the timescale of the functional connection. The true distributions of variables *X*_*t*_ and *Y*_*t*_ (e.g. neuronal signals) are not known a priori and must be estimated from repeated sampling of *X*_*t*_ and *Y*_*t*_ over multiple trials, as well as over several consecutive time bins if the time scale of the underlying signal is sufficiently slow. In general, the computation of robust approximations of information-theoretic quantities such as MI and TE from sample data is not trivial. We have used the library IDTxl (*30*) to compute TE for our data. BTE has two major advantages over naive measures of functional connectivity such as cross-correlation. First, it can handle arbitrary nonlinear relationships provided sufficient data are available and, second, it explicitly eliminates any spurious connections that are already well-explained by the past of the target variable. However, it only considers two variables at a time, and thus it is overly liberal when addressing higher-order interactions such as causal chains or common causes. As alternative to BTE, we therefore also used the Multivariate TE (MTE) metric, which only considers a functional connection existent if the estimated information transfer cannot be completely explained by the presence of any other connection. It also prioritizes interactions on short time scales over those further back in the past, in an attempt to deal with causal chains. The assumptions of MTE are unlikely to be fully met in the brain due to redundancy. The true functional connectivity thus is likely to be a compromise between that reported by BTE and MTE. In order to capture time-dependent changes in functional connectivity, we computed BTE and MTE across trial time in a sliding window of six 200-ms time bins.

### Validation of TE estimates using simulations

In order to gain intuition on the performance of the TE estimators, it is necessary to compare their accuracy with a ground truth model. For this purpose, we designed a model with known ground truth connectivity, generated dynamic data, estimated functional connectivity using BTE and MTE, and compared the estimates with ground truth (figs. S4 and S5). We validated connectivity estimates by computing binary connectivity matrices: a connection is present if p < 0.01, otherwise it is absent. In order to evaluate accuracy, we computed true positive (TP), true negative (TN), false positive (FP) and false negative (FN) rates, with ‘rate’ defined as the ratio of the number of connections of a given type, divided by the total number of possible connections *k·(k-1), w*here *k* is the number of regions considered.

We used a ‘chain model’ of 12 sequentially connected regions. Activity *X*_*i*_ in the *i*-th region was modeled considering a leak term, the weighted input from the previous neuron, and external input. By discretizing time in times bins of *dt* = 50 ms (20 Hz frame rate) we arrive at a linear discrete dynamical system

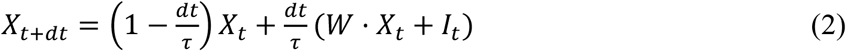

where *W*_*ij*_ = *w* ·δ(i+1, j) denote the connection weights and *I*_*t*_ the independent Gaussian noise input to each region. The connection weight *w* was set to one. The time scale for TE estimation was, dt=50 ms the time bin duration, *τ* = 200 ms is the time constant for the system dynamics. Each region received its own independent noise input which was propagated in the chain to the downstream regions. The chain model provides the known ground truth connectivity and, hence, is useful to evaluate the rates of FP (falsely identified connections) and FN (missed true connections) for a TE estimator.

In order to demonstrate the difference between BTE and MTE estimators, we present connection frequencies predicted by the model of sequentially connected regions and averaged over random initializations of the model. The connection value was taken to be the maximum over lags of 1 to 5-time bins. The more conservative MTE predicts only the strongest connections from one-time bin back (fig. S4), while cross-correlation and BTE predict chain jumps of 1, 2, 3 and 4 regions corresponding to lags 2, 3, 4 and 5 (fig. S4). While in this example the predictions of MTE appear to be true, in a more general scenario the same information could be carried by several connections with different lags. Thus, it is recommended to consider BTE and MTE as providing upper and lower bounds on true connectivity, respectively.

Using the chain model, we generated datasets of varying size (total number of data points), adhering to the structure of our experimental dataset. We have studied the effect of increasing the number of samples (time bins) while keeping other parameters fixed (fig. S4). We have also studied the effect of increasing the number of trials and found the results comparable (not shown). FP rate did not depend on data size; for BTE it fluctuated around 3-4% and for MTE it was below 1% (fig. S4). We also demonstrate that the absolute value of TE remained constant with varying data size for TP, but dropped for FN approaching the ground truth connectivity at infinite data size.

We also used the chain model to generate a fixed dataset with 400 trials, 12 regions, and 6 time bins, which corresponds to the length of the sliding window that we used for the analysis of experimental data. We normalized the dataset to its maximum value, and progressively added white noise. Both BTE and MTE broke down at a signal-to-noise ratio of approximately 0.3 (fig. S5). When we similarly ‘diluted’ our experimental data with noise, the drop out of connections had a similar rate as in the simulated curve, indicating a sufficiently high signal-to-noise ratio of the calcium signals. Finally, we used the chain model with the fixed dataset and then replaced a fraction of trials with white noise to simulate such effects as variance of timing and strategy across trials. Both BTE and MTE broke down (rate of TP dramatically decreased while FP increased) when the simulated connection frequency was reduced below approximately 0.2 (fig. S5). When we replaced a fraction of experimental trials with white noise (not shown), the TP and FP behavior was similar to the dynamics of the chain model MTE indicating that in our experimental dataset a trial-to-trial timing and strategy variability did not exceed 80%. Here we characterized TP and FP rates of the TE estimator against varying levels of noise. We attempted to model the effects of the instrument noise by varying the amplitude of the Gaussian noise added to each channel, while the effects of the biological noise (such as trial-to-trial variability in timing or the appearance frequency of the existing connection) were modeled by adding a variable number of ‘empty’ trials containing only Gaussian noise. However, designing a biologically plausible ground truth model to validate against the critical appearance frequency is difficult.

### Connection strength

The connectivity matrix for every session was calculated with the IDTxl (*30*) library. We defined a binary connection between each brain region based on significance level, i.e. if p ≤ 0.01 the connection was set to 1 (‘present’) and to zero otherwise (‘absent’). Connection strength was defined as the mean frequency of significant TE bins in a given analysis window (stimulus or action-reward), averaged across all mice and all expert sessions (n = 13 mice for Fig. 4 and n = 4 mice for Fig. 6). To estimate differences in functional connectivity between Hit and CR trials (Fig. 4H), we used a connectivity matrix inferred from MTE. For every mouse we averaged across all expert sessions. The resulting distributions of connection strength across mice for Hit and CR trials were tested with the Wilcoxon signed-rank test at every bin of trial time. Connections with p ≤ 0.05 were shown as significantly different for Hit vs CR trials.

### Analysis of functional network stability across-sessions

In order to evaluate the stability of emerging functional connections, we designed a test based on the probability distribution for the number of the functional connections shared between subsequent sessions. The number of connections observed in any given session is computed by summing up the entries of the corresponding binary connectivity matrix *M*.

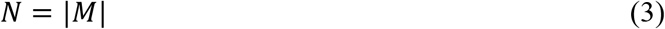

Note that the diagonal values of *M* are zero because, by definition, FC only includes connections between different channels. Thus, the highest possible number of connections is given by the number of off-diagonal entries of the matrix *M*, namely

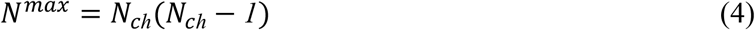

where *N*_*ch*_is the number of rows of that matrix (that is, the number of recorded brain regions).

The number of shared connections *N*_*sh*_ is defined as the sum of entries of the overlap between the matrices of two subsequent sessions. Denoting the “previous” session with *x* and the “next” session with *y*,

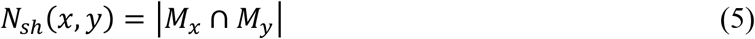

It can be easily seen that the maximum number of shared connections is given by the number of connections of the session that has the smallest of them

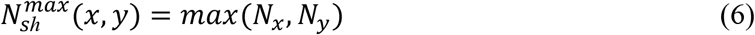

Given prior knowledge of *N*_*x*_and *N*_*y*_, the constraints on possible values of *N*_*sh*_can be summarized in the following: (1) *N*_*sh*_ ≥ *0* is non-negative; (2) *N*_*sh*_ ≤ *N*^*max*^ does not exceed the maximum number of possible connections; (3) 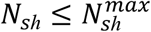 does not exceed the number of connections lowest among the two matrices; (4) 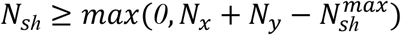 given sufficiently large *N*_*x*_and *N*_*y*_, there is a guaranteed non-zero overlap.

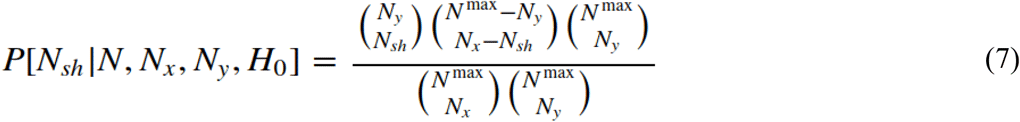

We construct the following null hypothesis *H*_0_: For all sessions *t* the functional connectivity *M*_*t*_ is completely random, except for the total number of connections *N*_*t*_. Namely, under *H*_0_ all permutations of *N*_*t*_ connections among *N*^*max*^possible connections are equiprobable. We proceed to calculate the probability distribution of the number of shared connections *N*_*sh*_ under *H*_0_, which will be used to test the significance of shared connections observed in real data.

The denominator is given by the number of possible permutations of *M*_*x*_ and *M*_*y*_. The numerator is constructed by first choosing *N*_*y*_ from *N*^*max*^, then choosing *N*_*sh*_ from *N*_*y*_, and finally distributing the non-shared connections of *M*_*x*_ among the remaining zeros of *M*_*y*_

We use the above distribution to calculate the p-values of the following two-sided tests: (1) 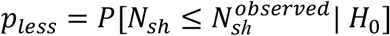 - the probability of getting as few of fewer shared connections as observed; (2) 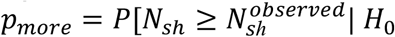 - the probability of getting as many or more shared connections as observed. Calculating the above p-values for observed data demonstrates that the number of the observed shared connections is consistently very unlikely under *H*_0_, proving that *H*_0_ does not hold for the observed data. This implies that the overlap between subsequent sessions is significantly different from a purely random one, mostly because there are many more shared connections than expected under *H*_0_.

Further, we use *S* =*log p*_more_as a measure of stability of the network. The larger the absolute value of *S*, the more shared connections there are, and the more stable is the network. We plotted *S* as a scatter plot of mouse performance in order to demonstrate the relationship between the network stability and learning. The scatter plots combined the results of all consecutive session pairs from all mice. We also performed a Mann–Whitney U test to determine whether the networks were more stable for expert or naïve mice. We visualized the results of this test using violin plots. Empirical distributions of *S* are highly asymmetric, and thus are difficult to accurately represent using bar plots. Finally, we used a sliding window to construct a trendline for the scatter plot to show the dynamic changes of stability with learning. For any given window, the likelihood of observing all data points in that window is given by

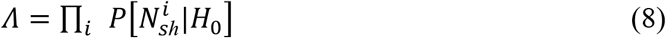

and the associated log-likelihood

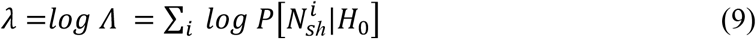

where *i* was used to index the data points within the sliding window. The log-likelihood is a linear function of the number of data points in the window *N*_*w*_, so we normalized the log-likelihood with that number to account for the fact that the number of datapoints in the window may vary. Thus, we defined the trendline as the arithmetic mean of the logarithms of individual probabilities, which translates to the geometric mean of the probabilities themselves.

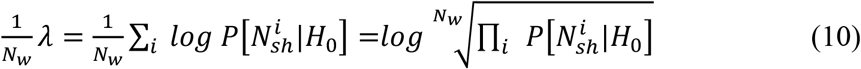

In order to smoothen the trendline, we replaced the window-based estimation of running average with kernel smoothing. We used a gaussian kernel with σ^2^ = 0.005 units of performance squared.

### Clustering coefficient

We used a local clustering coefficient as a measure of local change in connectivity during learning. For every region of the connectivity matrix estimated from the MTE analysis we calculated the number of outgoing connections *n*_out_. If *n*_out_ ≥ 2 and both target regions were also connected, we calculated the clustering coefficient as

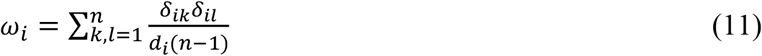

where we summed across all outgoing connections for the source *i* to pairs of targets (*k*, *l*) and normalize it by the node degree and number of targeted regions minus one. If we compared the changes in clustering coefficient for the stimulus window and the action-reward window, the connectivity matrix is first averaged across all connections within the given time interval and then the weighted clustering coefficient was calculated, where *δ*_*ik*_*δ*_*il*_ represent averaged across the time interval connection strength. Next, clustering coefficient within each time interval is averaged across respective groups for naïve and expert sessions (Fig. 4B) or for the groups for Hit and CR trials (Fig. 5B). Group differences are estimated with Mann–Whitney U test across mice.

## SUPPLEMENTARY NOTES

### Interpretation of the trial related brain activity

During the task execution we observed two prominent peaks of activity. The early peak is associated with the texture touch and decision related signals. The late peak likely reflects various components, including licking actions, responses to second auditory cue signaling the decision report phase, and reward- and punishment-related activity. Interestingly, early in training, when the reward was novel, the peak of calcium signals was higher for the reward itself (fig. S2). As mice subsequently started to associate reward to the available cues (presence of a texture and the auditory cue at 5 s signaling the report time) the onset of the late peak shifted to these cues and preparatory lick action. While the early peak was slowly established on the course of learning.

Because regional activities can be represented by mixtures of behavioral components, for our analysis we wanted to exclude the movement related activity. Such activity is present in the ‘go’ (Hit and FA) trials due to licking and additional body movement. We used two controls. First, we trained LDA (randomly selected 80% training data; 20% test data; 20-fold cross-validated) to classify task related information from the regional mean ΔF/F activity before the first lick, *i.e.* given by the early stimulus window from 3 - 3.5 s. Second, to control whether Hit/CR classification can be induced by action-related behavior, we trained LDA to classify action-related information for FA/CR trials as well in early stimulus window (fig. S3). Action-related classification accuracy peaked during pre-learning and first learning sessions and gradually decreased back to the chance level for late expert sessions. Most prominently a transient change during learning was expressed in CA1Py and M1 (p=0.02 and p=0.04 Mann–Whitney U test, fig. S3). Taken together results from (Fig. 2D and E and fig. S3) suggest that the increase of task related classification accuracy in early stimulus window during learning cannot be fully explained by the motor related changes in calcium signals.

---

**Fig. S1:**
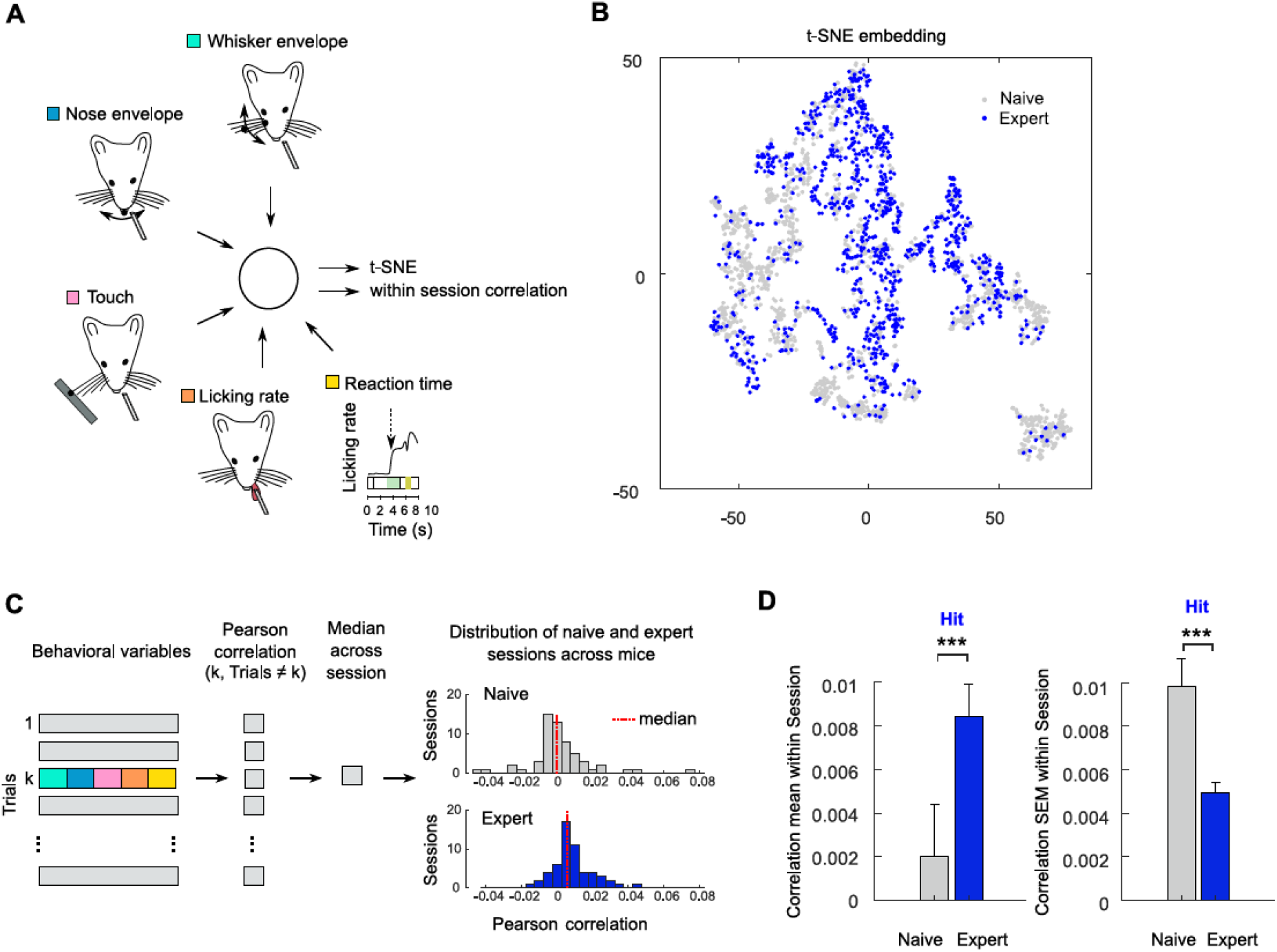
Behavior becomes more stereotypic during learning the texture discrimination task. (**A**) Behavioral parameters used in t-SNE embedding and for the calculation of the trial-to-trial Pearson correlation. (**B**) Example scatter plot for the t-SNE embedding of behavioral variables during Hit trials for one mouse. Whisker envelope, nose envelope, and touch were averaged during time interval of the texture approach and early texture presentation (2.5 - 3.5 s trial time), while licking rate was averaged across the whole trial time. All behavioral variables were scaled, that is z-scored (subtracted mean and divided by s.d. calculated for each behavioral parameter across all naïve and expert sessions for every mouse). Grey and blue dots represent separate trials during naïve and expert sessions respectively. (**C**) To quantify increase of stereotyped behavior with learning we computed trial-to-trial Pearson correlation coefficient. After calculating the median for each session and the mouse we sorted all sessions into naïve and expert groups. (**D**) Correlation across behavioral parameters increased during expert task performance. Bar plot showing mean ± s.e.m. (of median correlation coefficients for all mice and sessions; Mann– Whitney U test, *** p<0.001 for Hit). While the variability quantified by the s.e.m decreased for expert sessions (Mann–Whitney U test, *** p<0.001 for Hit). Learning related increase in trial-to-trial correlation and decrease in trial-to-trial variability of behaviorally relevant parameters suggests a shift from exploration to exploitation facilitating stable task performance and maximizing the amount of received reward in a given time of the session.

**Fig. S2:**
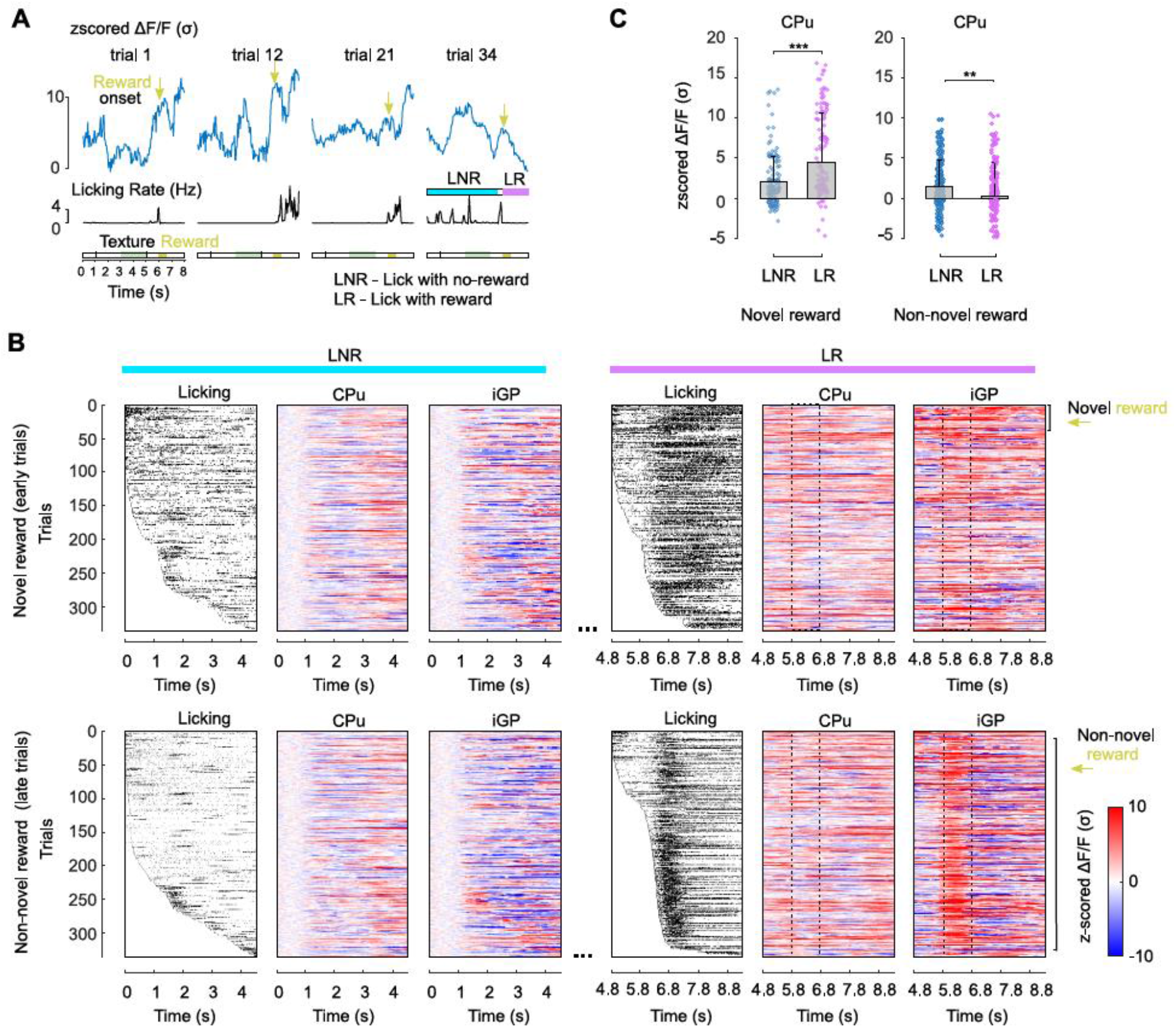
In naïve mice activity in CPu and iGP represent a mixture of licking action and reward. (**A**) Example of the z-scored ΔF/F calcium dynamics (s.d. and mean for each trial in the window 0 −1 s) recorded in CPu (blue) and the licking rate (black). Time for the water valve opening are shown with orange arrow. Texture presentation and reward time windows are shown with bars above the trial time. Magenta and cyan bars below trial 34 show periods for licking without reward (‘lick-no-reward’, LNR) and licking with reward (LR) for further comparison. (**B**) To illustrate the dependence of activity on lick and reward the trial time was separated into early (0 – 4.6 s) and late (4.8 – 9.3 s). Trials were sorted by the licking onset (from early to late lick initiation time) and corresponding z-scored calcium signals in CPu and iGP for LNR and LR conditions. Dashed bar (5.8 – 6.8 s) shows the time of water valve opening triggered by licking on the spout. (**C**) Bar plots show mean of z-scored activity + s.e.m. for the LNR condition (taken from 0-5.5 seconds of a trial time) and for the LR condition (5.8-8 seconds) in one mouse. We compare LNR and LR conditions for the early behavioral training when the reward was novel (first exposures to the reward) and for the later training when mice learned to lick for the reward (Mann–Whitney U test, ** p < 0.01, *** p < 0.001). CPu activity was higher for LR as compared to LNR when the reward was novel. However, activity during LR has gradually decreased with training.

**Fig. S3:**
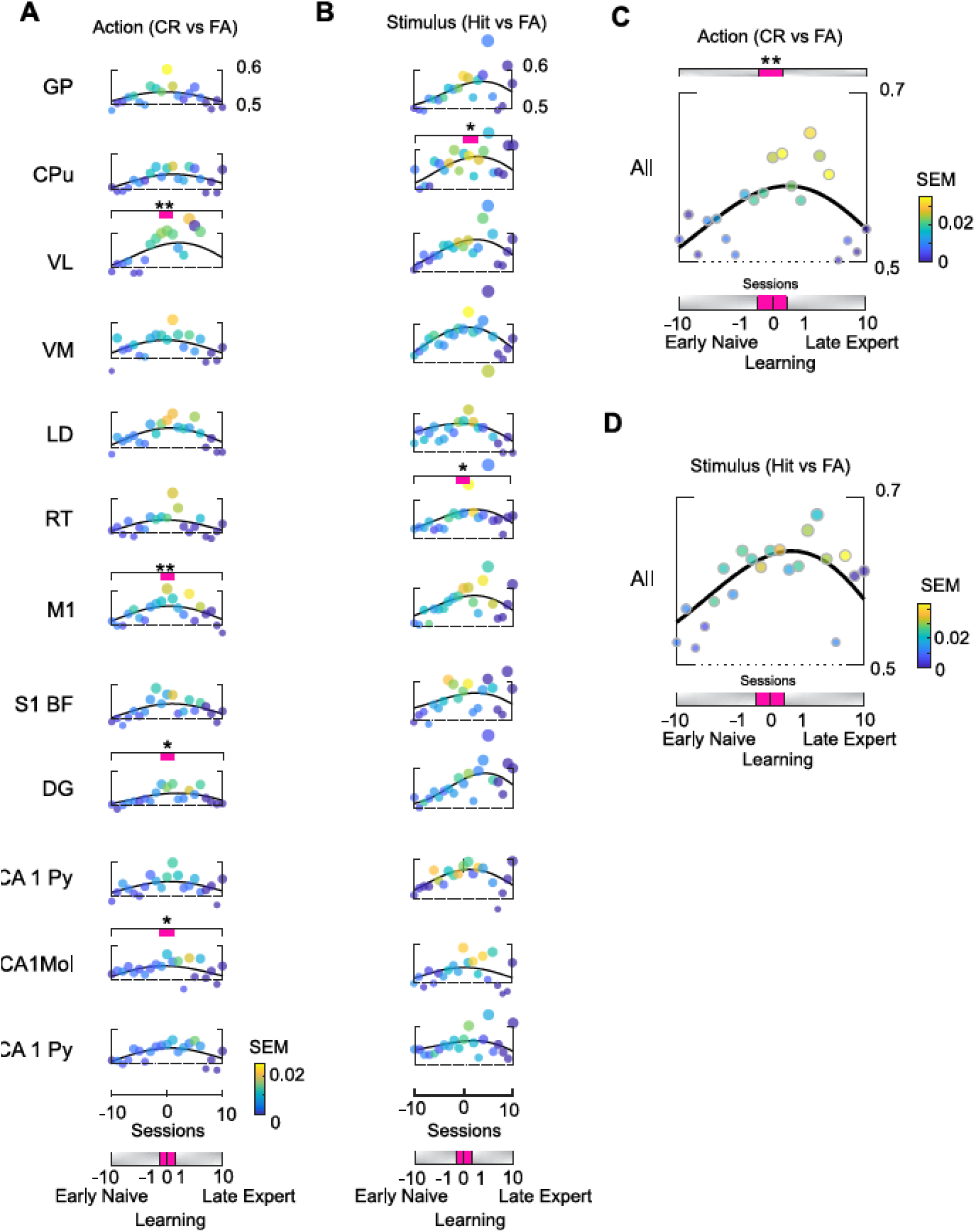
Action and stimulus predictive power appears in the population calcium dynamics during learning sessions. (**A**) Classification accuracy for action (CR vs. FA) averaged across n=14 mice and plotted versus training sessions (dashed horizontal line at 0 marks the pre-learning session); shown for 12 brain regions. Horizontal ticks represent 50 to 60% classification accuracy. Trial classifications were made for the LDA trained and tested for the calcium activity during the early texture presentation period (3 - 3.5 s). Color bar shows the s.e.m. Magenta bars above VL, M1, DG and CA1Mol show sessions which were used to compare trial discrimination accuracy across (Mann–Whitney U test n=14 mice, p=0.008 VL, p=0.003 M1, p=0.017 DG and p= 0.015 CA1Mol). (**B**) The same as **A** but for stimulus (Hit vs. FA) discrimination (Mann–Whitney U test n=14 mice, p=0.0013 for CPu, p=0.039 for RT). (**C**) Same as **A** but for all brain regions taken simultaneously for the LDA decoding. (**D**) Same as **B** but for all brain regions taken simultaneously for the LDA decoding.

**Fig. S4:**
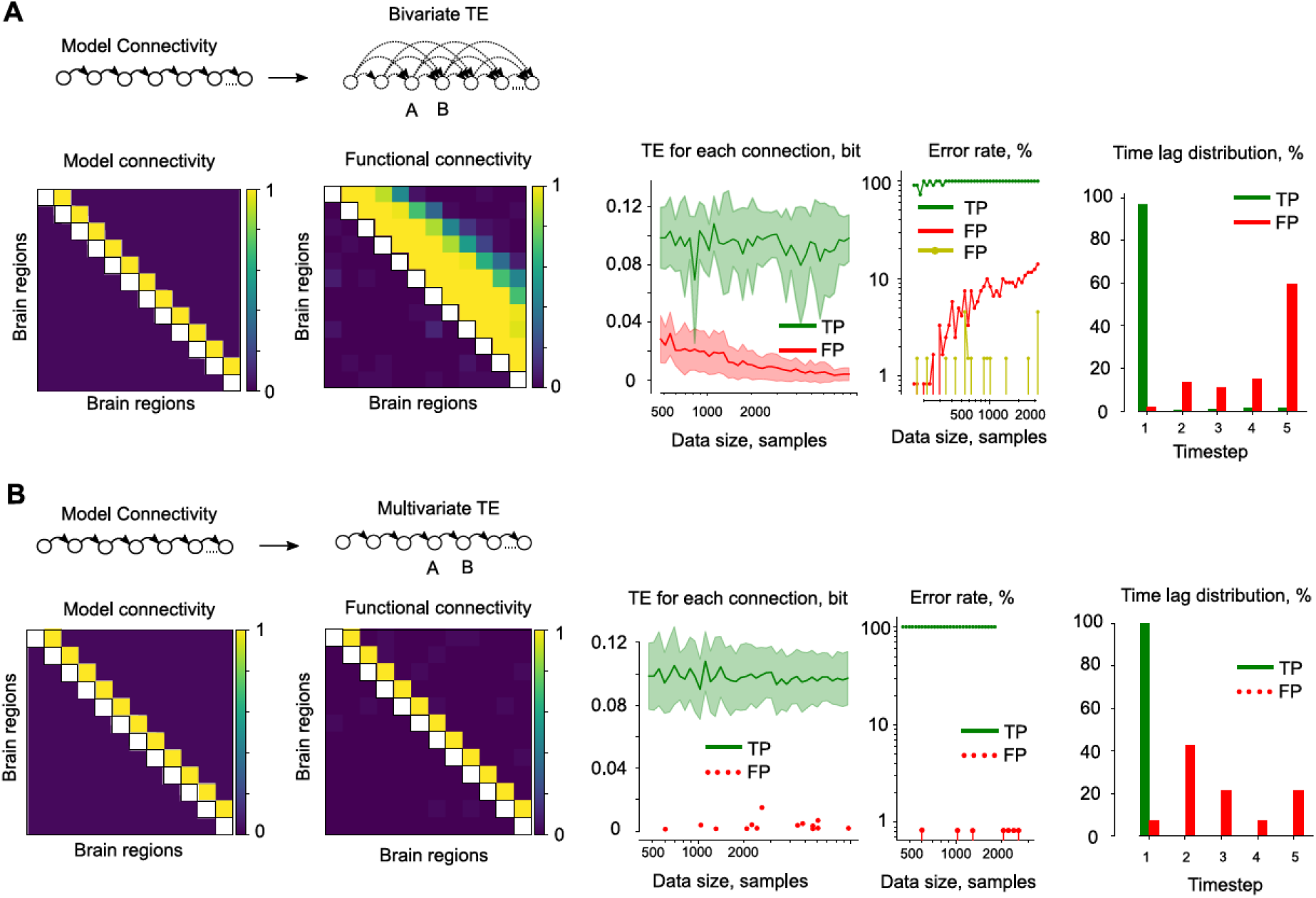
False positive connections are negligibly small as estimated by the model of sequentially connected regions. (**A**) Bivariate TE analysis on the model of sequentially connected regions. Left to right: model connectivity matrix representing ground truth connectivity (main diagonal has NaN values), estimated connectivity, TE for each connection estimated for the different data size, error rate (green: TP-true positive rate, red: FP-false positive rate for connections above the main diagonal, yellow: FP rate for connections below main diagonal), time lag estimated for true positive and false positive connections. Please note that bivariate analysis discovers connections coming from other source regions in the model of sequentially connected regions as the time lag is increasing (shown as additional diagonal rows 2, 3… etc. above the main diagonal on the connectivity matrix). Hence the conservative estimate of the false positive rate (red) is at 10% for the data size of approximately 1000 samples; however, at the time lag of 1 it is well bound to the margin of 2%. Less conservative estimate of the false positive rate (FP yellow) is below 2% when only connections below the main diagonal are considered. (**B**) Multivariate TE analysis on the model of sequentially connected regions. Left to right: model connectivity matrix representing ground truth connectivity (main diagonal has NaN values), estimated connectivity, TE for each connection estimated for the different data size, error rate, time lag estimated for TP and FP connections. As the data size is increasing the TE of the TP stays within the initial range of variance however TE of FP is decreasing in case of both bivariate and multivariate analysis. Multivariate TE is a conservative measure of the connectivity matrix because it eliminates connections at later time lags than 1 (because these connections can be often explained by a more recent information transfer).

**Fig. S5:**
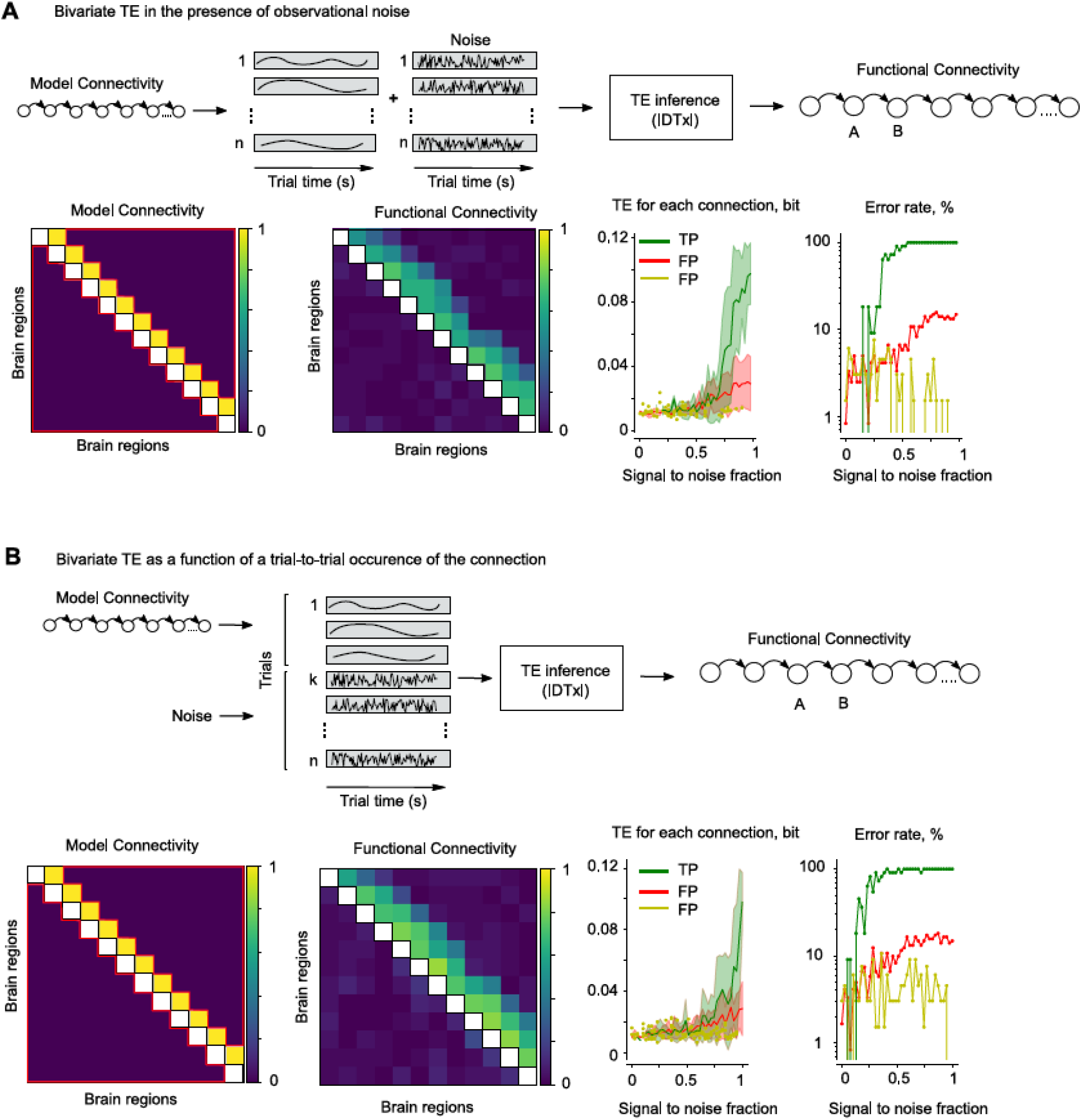
Loss of true connections happens when the noise starts to dominate the signal. (**A**) Bivariate TE analysis in the presence of observation noise. Left to right: model connectivity matrix representing ground truth connectivity (main diagonal has NaN values), estimated functional connectivity, TE for each connection estimated for different fractions of signal to noise (green: TP-true positive TE, red: FP-false positive TE for connections above the main diagonal, yellow: FP TE for connections below the main diagonal), error rate dependence on the fraction of signal to noise. The observation noise is simulated by adding Gaussian noise of different amplitudes to each trial with modeled temporal dynamics. TP are lost (green line) when the noise fraction is above 0.6. (**B**) Bivariate TE analysis as a function of trial-to-trial occurrence of the connection. Left to right: model connectivity matrix representing ground truth connectivity, estimated connectivity, TE for each connection estimated for different signal to noise occurrence frequency and the error rate dependency on the occurrence frequency. Trial-to-trial occurrence of the connection is simulated by varying proportion of modeled temporal dynamics to the trials containing only Gaussian noise (for example *k= n/2* corresponds to the signal to noise fraction of 0.5). TP are lost when the signal to noise fraction is less than 0.2.

**Fig. S6:**
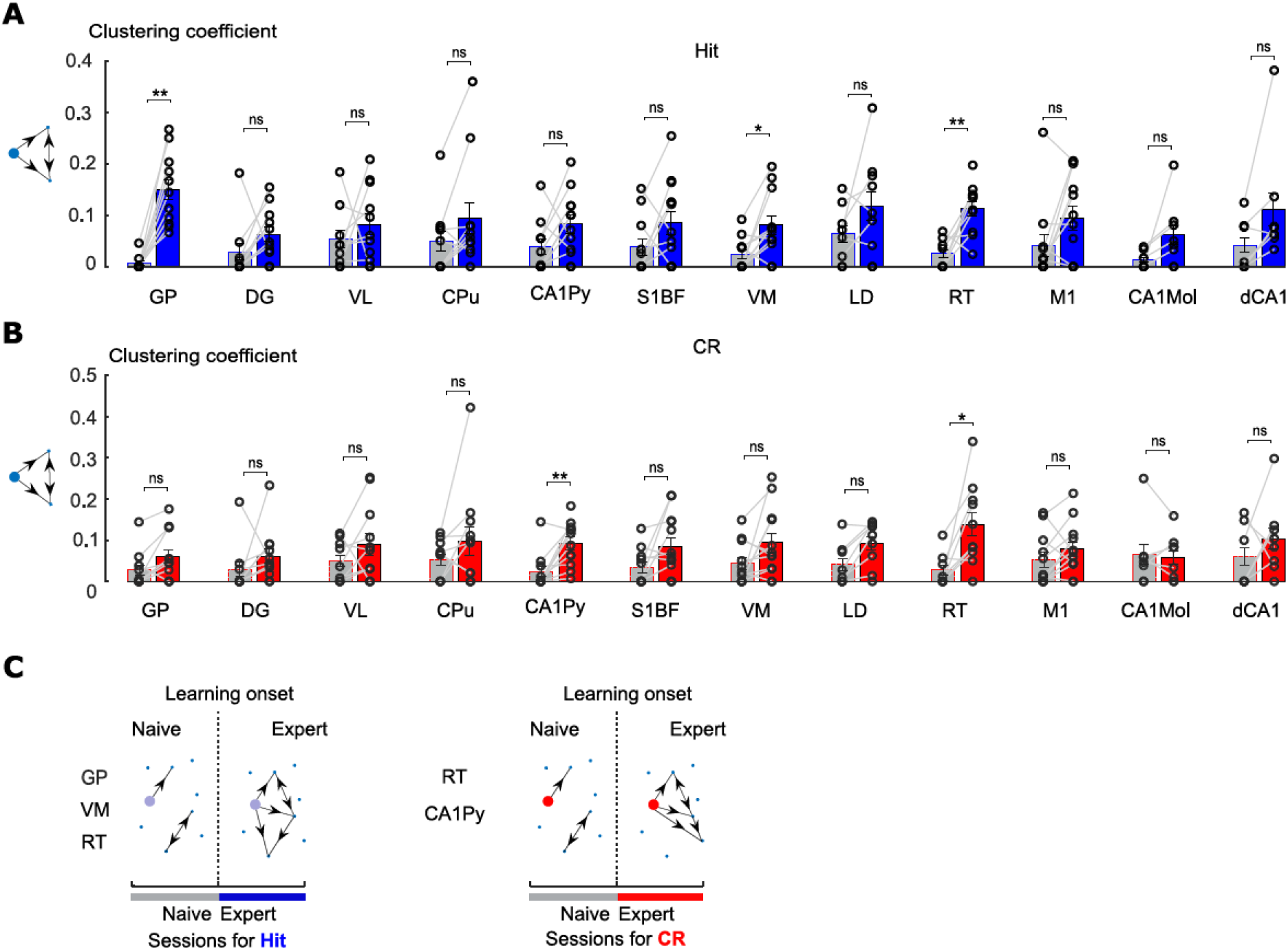
Clustering coefficient for all regions from the 12 channel network. (**A**) Bar plot comparing clustering coefficient for every region for naïve and expert phase (mean ± s.e.m. n=13 mice) for Hit sessions. (**B**) Same as **A** but for CR sessions. (**C**) Schematic illustration for the regions increasing their clustering coefficient

**Fig. S7:**
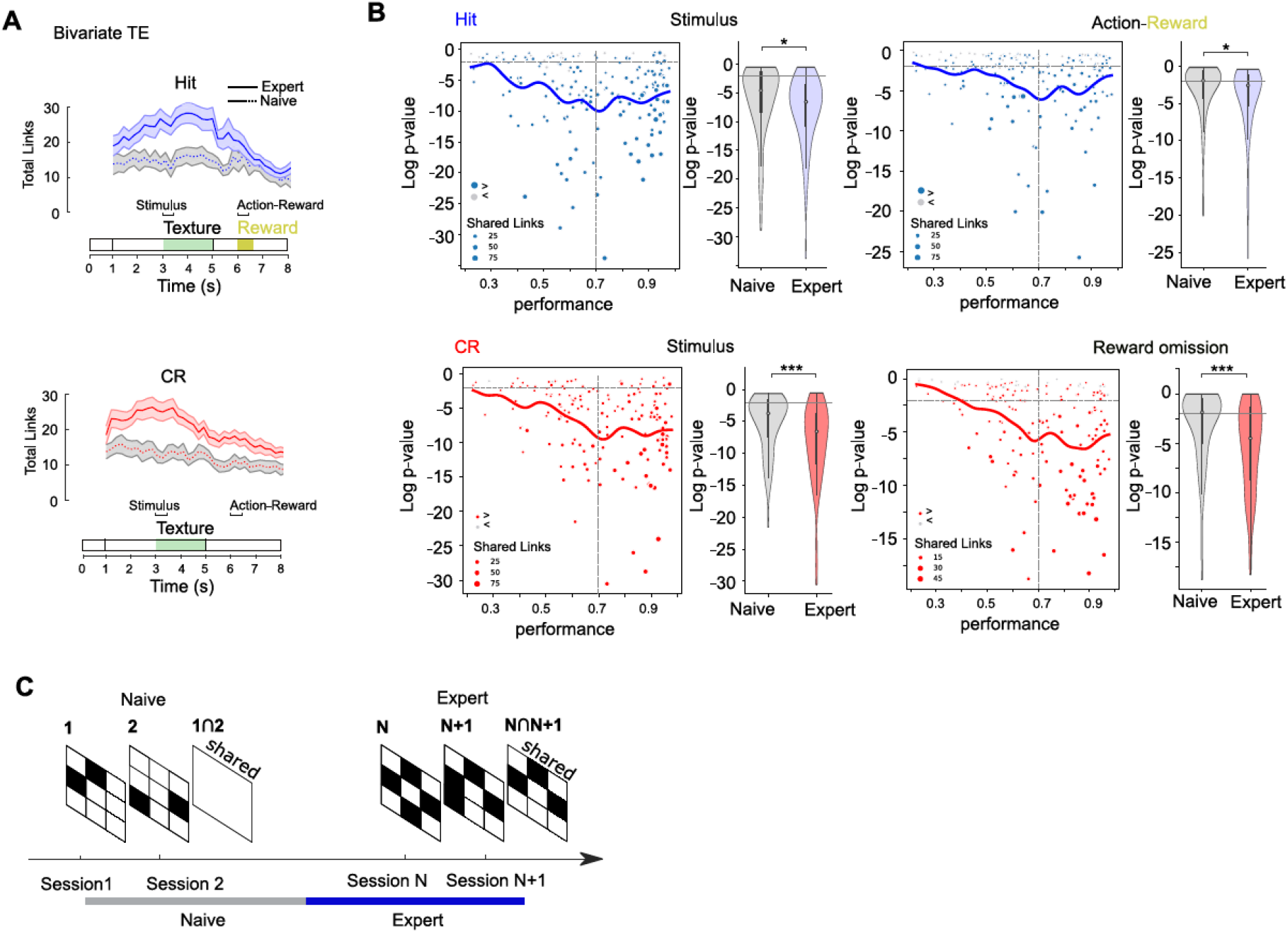
Functional network connectivity is increasing and stabilizing during learning also for the bivariate TE. (**A**) Total number of connections plotted as a function of the trial time for the bivariate TE (solid line and shaded areas represent the mean ± s.e.m. for the Hit and CR networks (blue and red respectively; mean± s.e.m. for the naïve mice is shown with dotted line and grey s.e.m.). (**B**) Scatter plot shows the Log(p) value to observe the same number of shared across neighboring sessions connections for the early texture presentation period versus the task performance. Circles are separate sessions for Hit (blue) and CR (red) trials. Solid line shows the regression to the scattered data. Violin plot on the right side shows Log(p) distributions for the naïve and expert phase (with the Mann–Whitney U test for the differences in naïve and expert phase network Log(p) values). (**C**) Same as B but for the lick-reward period. (**D**) Schematic illustration for the increase in number of shared connections during learning.

**Fig. S8:**
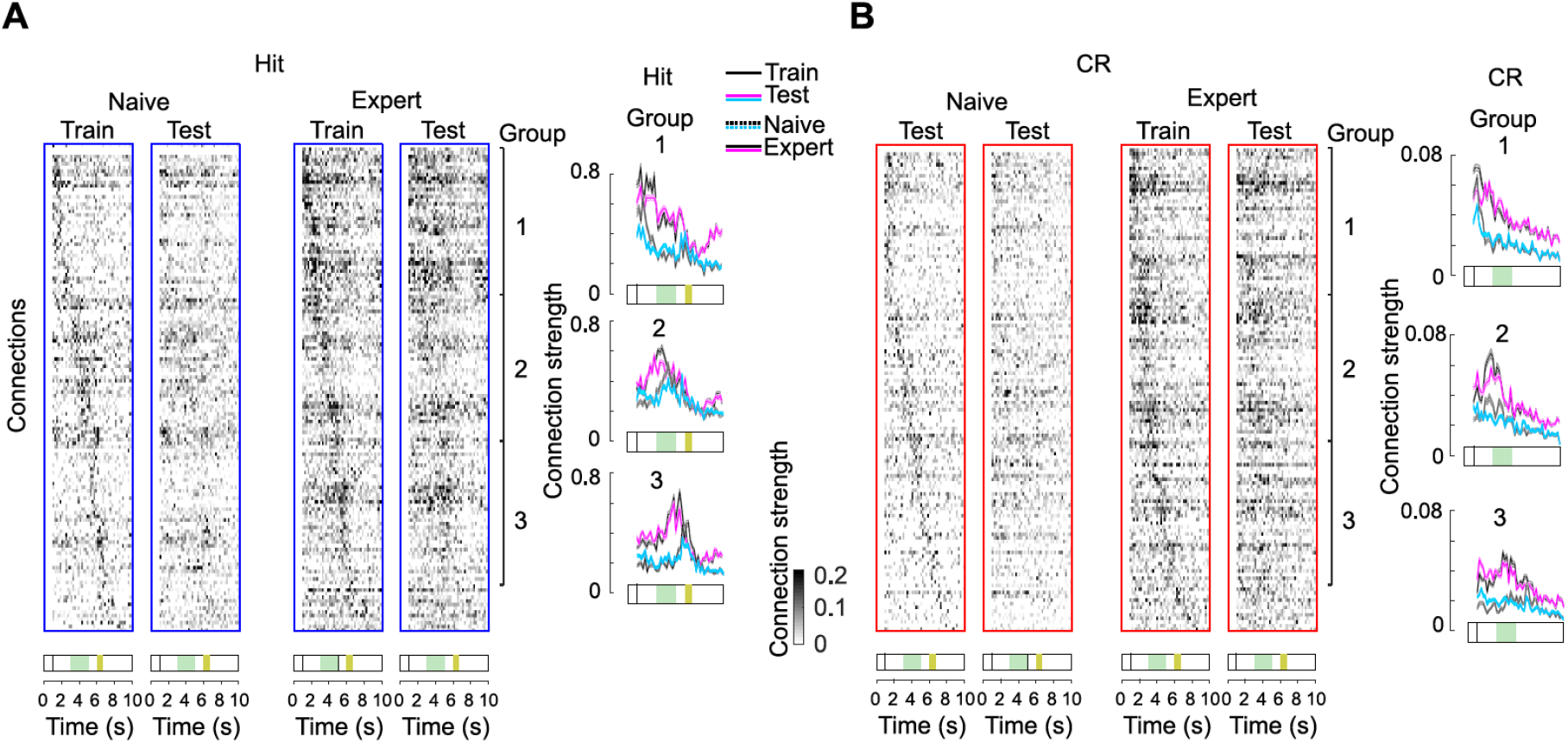
Segregation of functional connections into cue-, stimulus- and action-reward related groups. (**A**) Connection strength across trial time for naïve and expert phase. Dataset was randomly split to train (n=7 mice) and test (n=6 mice). Expert Hit sessions were sorted according to the peak trial time of each connection in the train dataset. We applied sorting indices from the train to the test dataset and plotted it for naïve and expert phases. We bootstrapped 20 times across 13 mice to verify that for different combinations of mice in train and test dataset connections were partially conserved within each of three major groups (1-40 cue-related, 41-80 stimulus-related and 81-120 action-reward-related). Temporal dynamics within each groups of connections for the train (mean ± s.e.m. across each group: black, dashed for naïve and solid line for the expert phase) and test (mean ± s.e.m. across each group: magenta, dashed for naïve and solid line for the expert phase). Mean for train and test overlapped indicating consistent connection assignment across mice to either cue-, stimulus- or action-reward group. Furthermore, we observed both strengthening of connections with learning and a similar to the ΔF/F shift of connection recruitment to the earlier times of the trial (comparison of dashed and solid curves particularly for stimulus- or action-reward groups). (**B**) Same as A but for CR sessions.

**Fig. S9:**
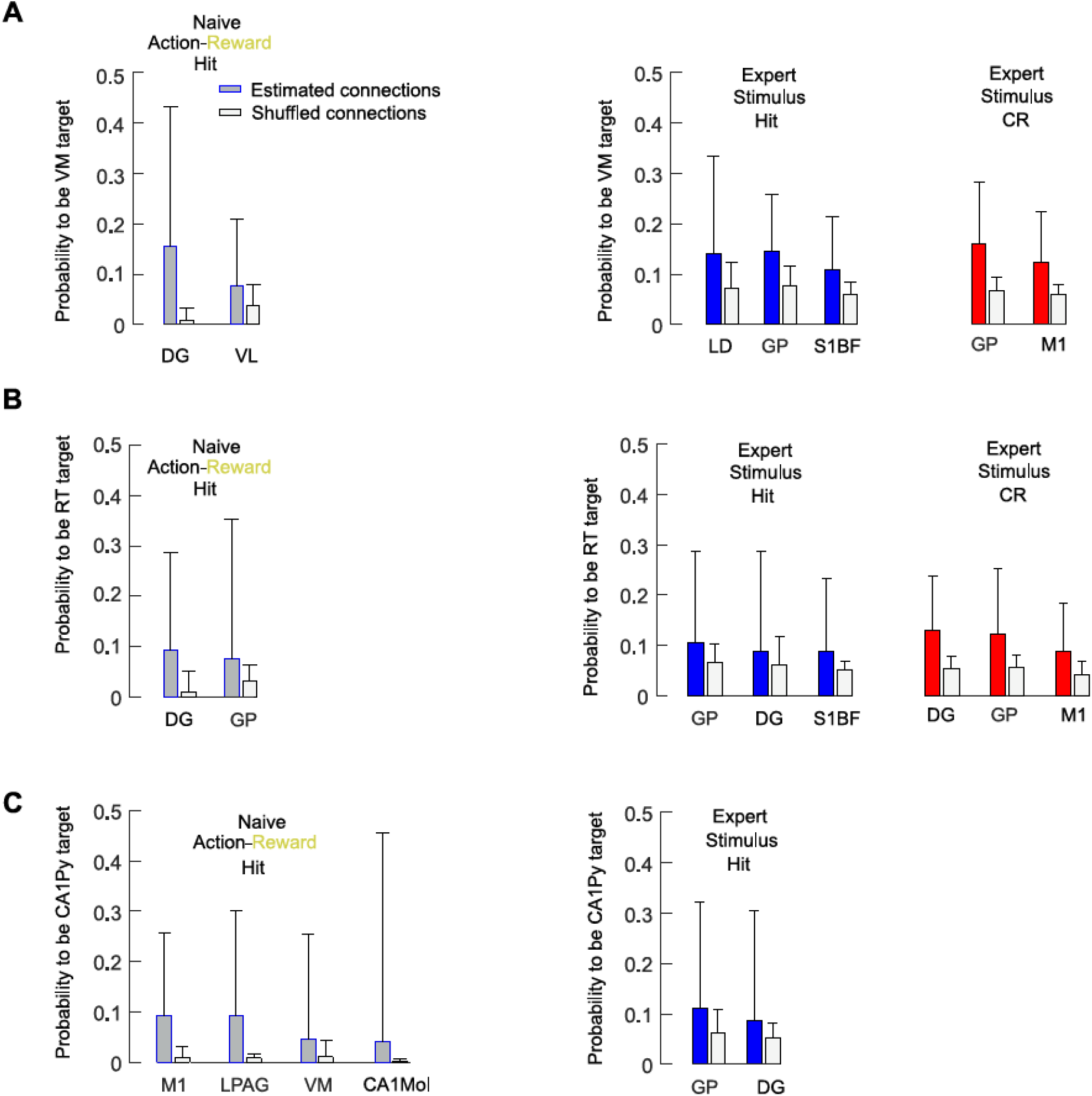
Target regions of VM, RT and CA1Py. Similar to iGP, VM, RT, CA1Py increased clustering coefficient during learning. (**A**) Probability to be targeted by VM (mean + 3 s.d., Hit blue and CR red) comparing action-reward for the naive phase and early stimulus presentation period for expert phase. Light grey bars show mean + 3 s.d. of probability to be targeted by VM for shuffled connections. Only target regions above shuffle mean + 3 s.d. are shown. (**B**) Same as **A** but for target regions of RT. (**C**) Same as **A** but for target regions of CA1Py.

**Fig. S10:**
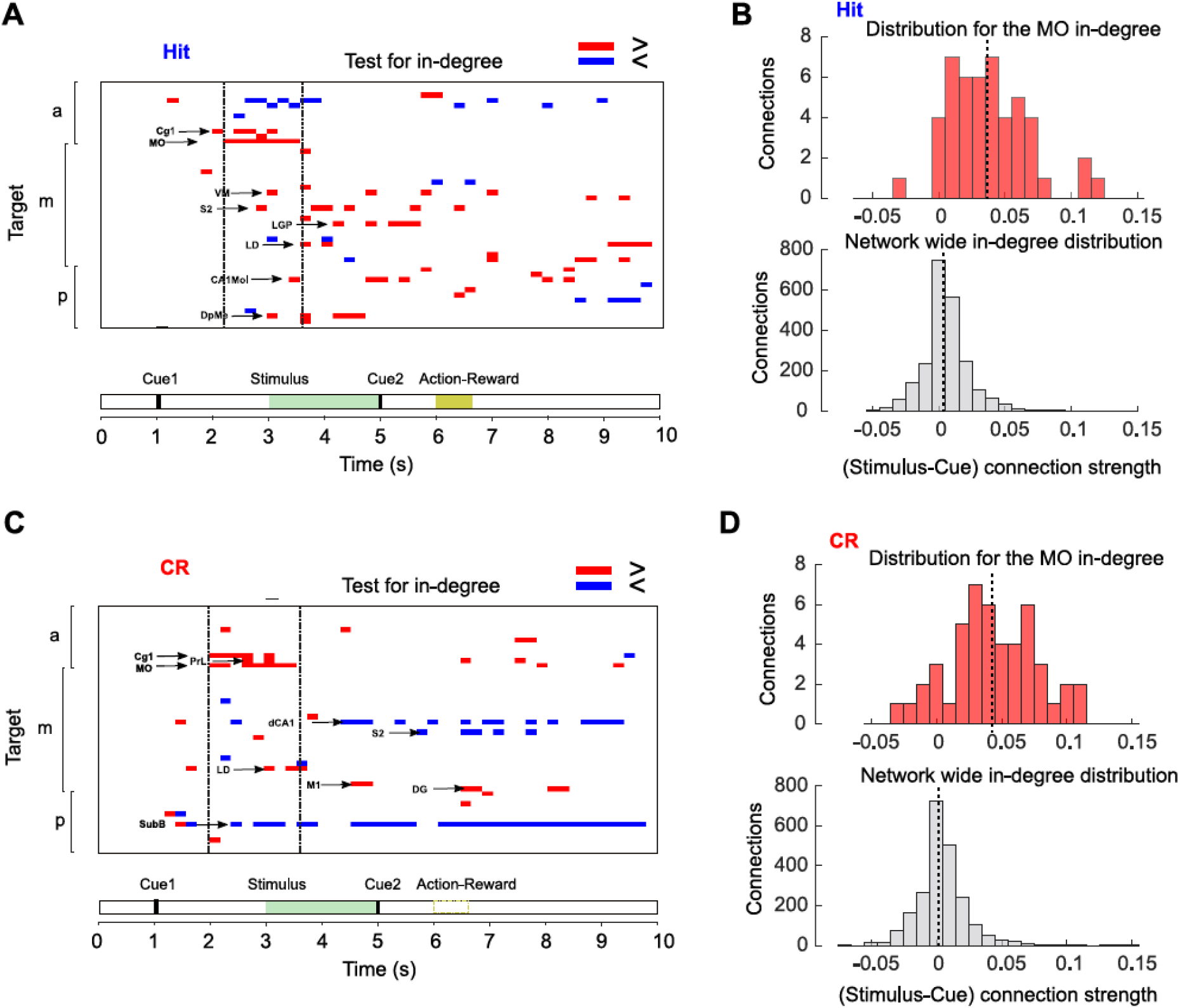
Dynamic shift of the input to prefrontal and orbitofrontal cortical regions during texture approach and early texture discrimination period. (**A**) Plot for the regions changing their in-degree distribution during Hit trials. For each region we tested if the median of the in-coming connection strength was significantly different as compared to the in-degree distribution for rest of the network (Mann–Whitney U test for the averaged across 4 mice connectivity matrix with subtracted connection strength during the cue period, p<1%; red: if regional median connectivity increased; blue: if decreased). (**B**) Example of in-degree distribution during the early stimulus period (3 – 3.6s) for the MO (top, red) as compared to the in-degree distribution for the rest of the network (bottom, grey). Dashed line shows the median for each distribution. (**C**, **D**) Same as **A** and **B** but for the CR trials.

**Supplementary Table 1.**
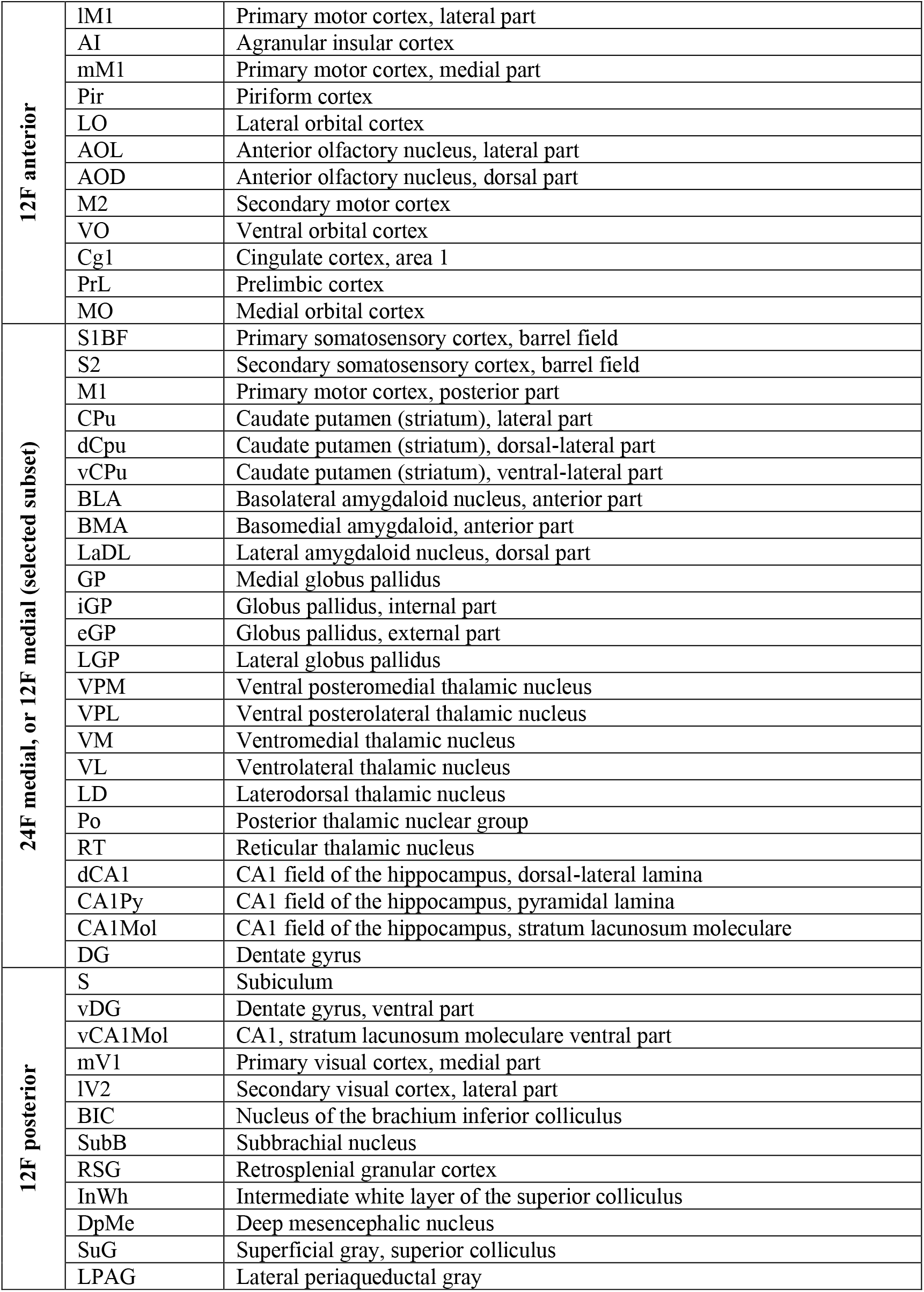
List of brain regions targeted in the multi-fiber photometry experiments. We mainly adhere to the nomenclature of Paxinos and Franklin. In a few cases we applied further subdivisions, e.g., for medial and lateral M1 and for sublaminae in CA1.

## Notes

### Competing Interest Statement

The authors have declared no competing interest.

